# USP50 suppresses alternative RecQ helicase use and deleterious DNA2 activity during replication

**DOI:** 10.1101/2024.01.10.574674

**Authors:** Hannah L. Mackay, Helen R. Stone, Katherine Ellis, George E. Ronson, Alexandra K. Walker, Katarzyna Starowicz, Alexander J. Garvin, Patrick van Eijk, Alina Vaitsiankova, Sobana Vijayendran, James F. Beesley, Eva Petermann, Eric J. Brown, Ruth M. Densham, Simon H. Reed, Felix Dobbs, Marco Saponaro, Joanna R. Morris

## Abstract

Mammalian DNA replication employs several RecQ DNA helicases to orchestrate the faithful duplication of genetic information. Helicase function is often coupled to the activity of specific nucleases, but how helicase and nuclease activities are co-directed is unclear. Here we identify the inactive ubiquitin-specific protease, USP50, as a ubiquitin-binding and chromatin-associated protein required for ongoing replication, fork restart, telomere maintenance and cellular survival during replicative stress. USP50 supports WRN:FEN1 at stalled replication forks, suppresses MUS81-dependent fork collapse and restricts double-strand DNA breaks at GC-rich sequences. Surprisingly we find that cells depleted for USP50 and recovering from a replication block exhibit increased DNA2 and RECQL4 foci and that the defects in ongoing replication, poor fork restart and increased fork collapse seen in these cells are mediated by DNA2, RECQL4 and RECQL5. These data define a novel ubiquitin-dependent pathway that promotes the balance of helicase: nuclease use at ongoing and stalled replication forks.

## Introduction

DNA replication is fundamental for genomic integrity. Obstacles to replication, including unrepaired DNA lesions or extensive secondary structure, can block the progression of replicative polymerases causing fork stalling, fork collapse, and generating DNA breaks ^1^. Hundreds of forks may stall during each S phase in a human cell, and the frequency increases in cells exposed to genotoxic or oncogenic stresses. Pathways to recover stalled and broken replication forks are utilised to resolve impediments to replication so that DNA synthesis can be completed. These pathways include reversal and stabilisation followed by restart; repriming; post-replicative repair; template switching; and double-strand break (DSB)-mediated recovery. Faults in processing obstacles or restoring replication following processing increase genomic instability leading to tumorigenesis ^2^.

RecQ helicases are a highly conserved family of helicases that have essential roles in replication and DNA repair^3^. They contain the core helicase domain (DEAD/DEAH box, helicase conserved C-terminal domain) and possess 3’ to 5’ unwinding directionality capable of unwinding a variety of structures; they can also anneal complementary ssDNA and perform branch migration (reviewed in ^4,5^). There are five human RecQ helicases: RECQL1, WRN, BLM, RECQL4, and RECQL5. Four are linked to human syndromes characterised by cancer predisposition and/or premature ageing: Werner’s syndrome (*WRN*), Bloom’s syndrome (*BLM*), and Rothmund-Thomson, RAPADILINO, and Baller-Gerold syndromes (*RECQL4*) ^4, 6^. Recently two families with a genome instability disorder named RECON syndrome have been found to carry biallelic mutations in *RECQL1*^7^. Mutations in all five helicase genes are associated with genomic instability and increased cancer risk^8^. The WRN helicase has been identified as a synthetic lethal target of cancers with high levels of microsatellite instability ^9, 10, 11, 12^. Its helicase activity is required to process cruciform structures formed of large (TA)_n_ repeats generated though microsatellite instability over time^13^.

Three of the RecQ helicases are employed in the restart of stalled replication forks. BLM deficient cells restart poorly after Aphidicolin or Hydroxyurea expossure^14^. RECQL1 restores stalled and reversed forks exposed to TOP1 inhibitors and to several other types of replication stress ^15, 16^. WRN facilitates the progression of stalled forks formed under normal physiological conditions or after exogenous genotoxic stress^17, 18, 19^. It has been implicated in the recovery of arrested forks^20, 21^, and was recently shown to contribute to the processing of stalled and reversed forks to promote restart ^22^. Additionally, the RECQL5 helicase may be used under certain circumstances. RECQL5 is recruited to stalled forks and can cooperate with WRN *in vitro*, its over-expression improves cell survival in the presence of the replication-blocking agent thymidine, and RECQL5 supports replication in WRN-deficient cells ^23, 24^. How RecQ helicases, which unwind similar DNA structures *in vitro* ^25, 26, 27, 28^, are deployed at different times and at different structures in cells is not clearly defined.

The human nuclease-helicase DNA2 recently emerged as critical to stalled replication fork processing (reviewed in ^29^), where it alleviates replication stress by promoting resection ^30^. DNA2 functions with BLM and WRN helicases in DNA end resection^31, 32^, but its role in replicative stress is linked to WRN, where DNA2:WRN degrade reversed forks to promote restart ^22^. DNA2 is vital to ongoing replication ^30, 33, 34, 35^ and biallelic *DNA2* mutations have been identified in patients with Seckel syndrome and primosmarcrdial dwarfism, conditions associated with under-replication^36, 37^.

Recently compound heterozygosity of *DNA2* mutations has been associated with severe growth failure and the clinical characteristics of Rothmund-Thomson syndrome ^38^, a condition previously linked to *RECQL4*. Thus, a critical question is how the relationship between DNA2 and particular RecQ helicases is promoted in particular contexts.

Ubiquitin (Ub) modification pathways are a central means to respond to and fine-tune replication fidelity. These modifications act in both the machinery of unperturbed replication and, most prominently, in the supporting pathways that tolerate, repair or respond to replication difficulties ^39^. Ub is a versatile protein acting both as a signal for protein turnover and providing a novel interaction face for the recruitment of factors that repair or tolerate replicative difficulties. Ub-interacting proteins carry one or more structurally diverse Ub-binding domain to drive such exchanges ^40^. Ub is conjugated to proteins through a three-enzyme cascade, whereas the processing of Ub from proteins acts to restrain the Ub signal. The role of Ub in replication is complex, and many of the pathways it regulates are poorly understood.

Here we expand our understanding of how RecQ helicases and DNA2 are controlled during replication. Our data indicates that USP50, an inactive member of the ubiquitin-specific-processing protease (USP) family of deubiquitinating enzymes, can bind Ub via its conserved USP domain. This ability of USP50 is critical for its localisation to chromatin, to nascent DNA and for the protein to promote normal replication kinetics. USP50 supports the ability of WRN to localise to stalled replication forks and interact with FEN1 and 9-1-1. USP50 promotes fork restart, suppresses MUS81-dependent breakage and promotes lagging-strand telomere stability. However, it is not vital to the survival of cells with high levels of microsatellite instability. Remarkably, cells lacking USP50 exhibit DNA2-, RECQL4-, and RECQL5-dependent ongoing replication and fork restart deficiencies. These findings link a novel Ub-binding protein to the correct coordination of RecQ helicases and DNA2 in replication.

## Results

### USP50 recruitment to chromatin is ubiquitin-dependent and is enriched at stalled replication forks

Two previous RNA interference screens have highlighted USP50 as potentially important to replication ^41, 42^. The USP class of deubiquitinating enzymes are cysteine proteases characterised by a catalytic domain divided into a series of conserved regions. Human USP50 lacks the conserved acidic residue of the catalytic triad and fails to process Ub-β -galactosidase^43^, and is classified as a non-protease homologue of USPs (uniprotkb/Q70EL3). The Alphafold USP50:Ub structure predicts a complex similar to structures reported for USP domains with Ub (e.g. PDB: 3n3k ^44^). We examined USP50 protein sequences from 24 diverse species. We noted 49 invariant and 39 highly conserved amino acids out of 339 total, suggesting the conservation of some aspects of the protein, including of the predicted Ub interaction face (Figure 1A, Supplementary Figure 4). The USP-family of deubiquitinating enzymes interact with the hydrophobic patch of Ub centred around isoleucine 44 (Ub: Leu8, Ile44, Val70) ^45^. In the predicted USP50:Ub structure, Ub Ile-44 is close to USP50 Ile-141 (Figure 1A), and leucine or isoleucine is found at this position in all 24 species. To test whether USP50 can bind Ub we expressed FLAG-USP50 and I141R-FLAG-USP50 mutant with Myc-tagged Ub and performed FLAG immunoprecipitation. We found that WT-USP50 co-precipitated high molecular weight Ub conjugates, but the mutant USP50 showed a reduced ability to co-purify Myc-Ub (Figure 1B), suggesting that USP50 binds Ub conjugates, at least in part through its predicted Ub-binding face.

**Figure 1.**
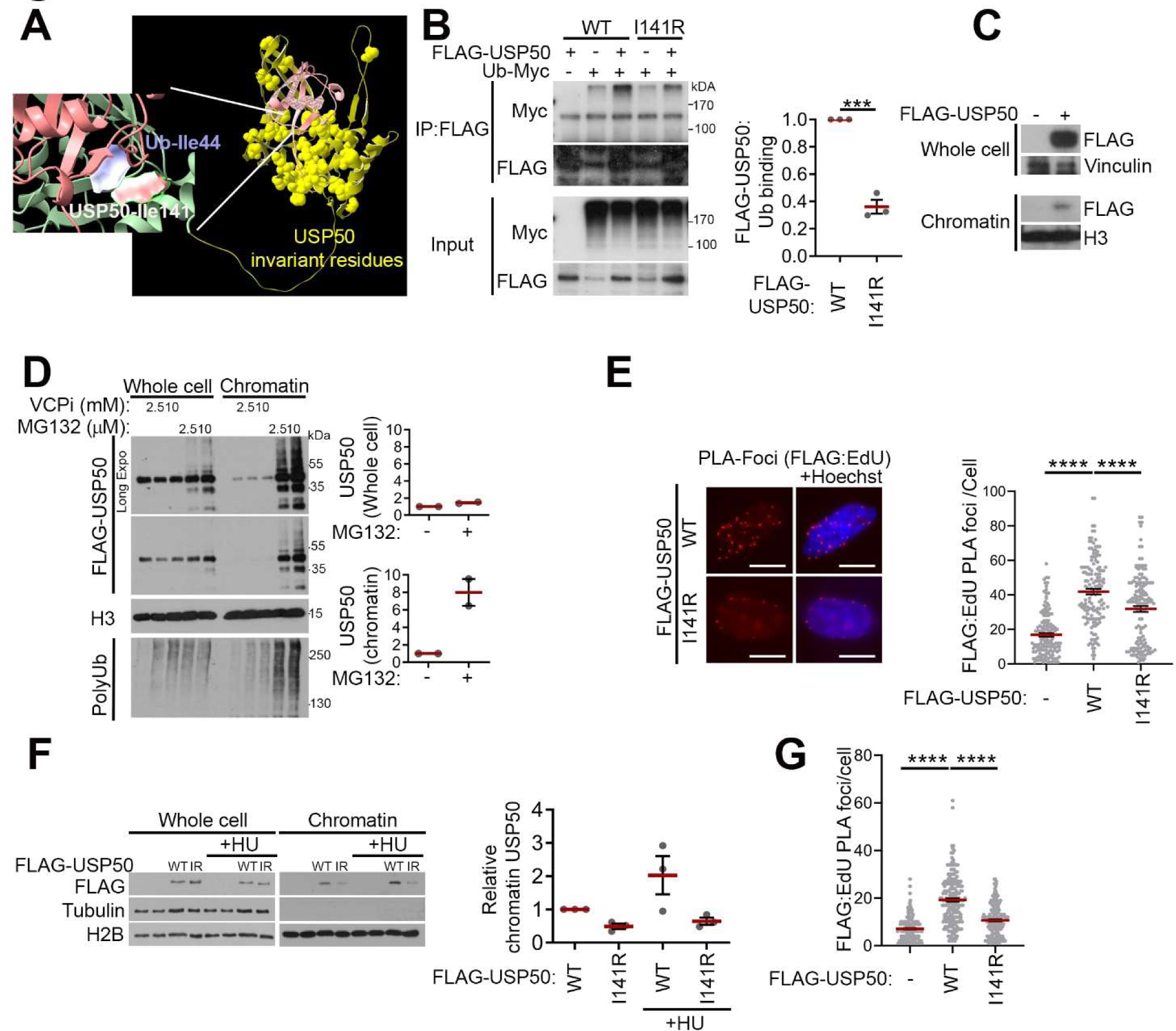
USP50 is recruited to chromatin in a ubiquitin-dependent manner and is enriched at stalled replication forks. A USP50:Ub interaction predicted by Alphafold2. Electrostatic densities of invariant USP50 residues are shown in yellow. In the inset Isoleucine-44 of Ub and isoleucine-141 of USP50 are shown as electrostatic density. B Immunoprecipitation of FLAG epitopes from HeLa cells expressing FLAG-USP50 or I141R-FLAG-USP50 and Myc-Ub, probed for FLAG and Myc (left) and quantification (right) of Myc-Ub from 3 independent experiments, and normalised to both Myc-Ub and FLAG-USP50 expression in the whole cell lysate. Red bars indicate mean, error bars are SEM. C Representative blot illustrating the presence of FLAG-USP50 in the whole cell lysate (Vinculin loading control) and chromatin enriched fraction (H3 loading control) following a chromatin fractionation assay. D Representative blot of whole cell lysate and the chromatin enriched fraction from HeLa cells expressing FLAG-USP50 and treated with VCPi, CB-5083, for 3 hours, or MG132 for 4 hours (left). Quantification (right) of FLAG-USP50 in the whole cell lysate normalised to H3 and chromatin fraction normalised to whole cell lysate levels of FLAG-USP50. Results are plotted relative to FLAG-USP50 in vehicle *Vs* 10 µM MG132 treated samples, from 2 independent experiments. Red bars indicate the mean, error bars are SEM. E FLAG-USP50 proximity to DNA labelled with EdU for 24 hours measured via PLA, using antibodies to FLAG and Biotin-EdU. HeLa cells with and without shUSP50 treatment and complementation with FLAG-USP50 or I141R FLAG-USP50 were scored for red PLA foci formation. Representative images (left) include scale bars of 10 µm. Quantification (right) of PLA foci formation per cell was determined by 3 independent experiments (n >150 cells per condition). Red bars indicate mean, error bars are SEM. F Representative blot (left) of whole-cell lysate and the chromatin enriched fraction from HeLa cells expressing FLAG-USP50 or I141R-FLAG-USP50, untreated or treated with 5 mM HU for 3 hours. Graph (right) shows quantification of FLAG-USP50siR in the chromatin fraction relative to FLAG-USP50siR in the untreated sample, from 3 independent experiments. Red bars indicate mean, error bars are SEM. G FLAG-USP50 proximity to DNA labelled with EdU for 15 min (followed by 3 hours 5mM HU treatment), measured via PLA, using antibodies to FLAG and Biotin-EdU. HeLa cells with shUSP50 treatment and complemented with FLAG-USP50 (first two conditions) or I141R FLAG-USP50 were scored for PLA foci. In the first condition, EdU was omitted. Quantification of PLA foci formation per cell was determined by 3 independent experiments (n >150 cells per condition). Red bars indicate mean, error bars are SEM.

If USP50 has a direct role in replication, it might be expected to be associated with chromatin. We fractioned cells expressing exogenous FLAG-USP50 and noted a proportion of USP50 co-fractionated with chromatin (Figure 1C). To address whether Ub can influence USP50 localisation, we treated cells with the proteasome inhibitor MG132 to reduce Ub conjugate turnover and the VCP/p97 inhibitor CB-5083 to reduce the extraction of Ub-conjugated proteins ^46^. Both treatments increased Ub conjugates in whole-cell lysates; proteasome inhibition enriched Ub conjugates within chromatin more than VCP inhibition and greatly increased the chromatin association of USP50 (Figure 1D). To further assess USP50 localisation, we incubated cells with the nucleotide analogue EdU for 24 hours to label DNA. We used a proximity ligation assay (PLA) with antibodies to the analogue (EdU) and to the FLAG fused to USP50. This method indicated an association of FLAG-USP50 with DNA and less association of the Ub binding mutant, I141R-FLAG-USP50 (Figure 1E).

To assess whether USP50 localisation relates to stalled replication, we examined USP50 co-fractionation with chromatin following hydroxyurea (HU) treatment. We observed a two-fold increase in FLAG-USP50 levels at chromatin, whereas the level of chromatin-associated I141R-FLAG-USP50 did not increase following HU exposure (Figure 1F). To address whether USP50 locates specifically to stalled replication forks, we labelled nascent DNA with a short (15-minute) pulse of EdU, then slowed replication with 3 hours of HU treatment^47^ and examined the proximity between FLAG-USP50 and EdU. This method showed enrichment of FLAG-USP50, and to a lesser extent I141R-FLAG-USP50 at EdU-labelled DNA (Figure 1G). Thus, USP50 localisation to chromatin and to nascent DNA is increased following HU treatment in a manner that largely depends on its Ub-binding face.

### USP50 promotes replication in unperturbed and stressed conditions

Human USP50 mRNA is part of cluster 71, defined as a testis-DNA repair cluster (confidence 0.99), and its expression is at low levels in most other tissues (^48^ and Human Protein Atlas, proteinatlas.org). Indeed, we could not detect USP50 in Hela cell lysates by immunoblot. As the screens suggesting an impact of targeting USP50 on replication were performed in Hela and A549 cells ^41, 42^, we wished to test whether USP50 protein is relevant to replication despite its low-level expression. We generated HeLa cells bearing an inducible shRNA to USP50, which we demonstrated depleted wild-type, exogenous FLAG-USP50 (Supplementary Figure 1A). We next generated FLAG-tagged USP50 resistant to that shRNA, which was integrated into an inducible site, expressed upon doxycycline treatment, named FLAG-USP50 hereafter. We made a second line in the same way with the exception that Ile-141 was mutated to arginine, named I141R-FLAG-USP50 hereafter (Supplementary Figure 1B). We then examined replication fork structures using the DNA fibre assay, incorporating two nucleotide analogues sequentially and recording the types of structures observed (illustrated in Supplementary Figure 1C). Strikingly, cells treated with USP50 shRNA exhibited increased first-label terminations, reduced ongoing forks and greater asymmetry between second labels from first-label origins (Figure 2A & B), indicating that USP50 is needed for ongoing replication. shUSP50-expressing cells complemented with FLAG-USP50 had replication features comparable to the control cells, whereas expression of I141R-FLAG-USP50 did not improve replication defects in shRNA-treated cells (Figure 2A & B). We examined the stability of forks stalled by HU treatment and observed that cells expressing shUSP50 had slightly shortened nascent DNA, suggesting a reduced ability to protect stalled structures (Supplementary Figure 1D). We next tested the ability of stalled replication forks to restart, employing an alternative version of the DNA fibre assay in which the second label is applied after washing out the HU. USP50 shRNA-treated cells exhibited a decrease in restarted forks and an increase in first-label terminations, which could be complemented by FLAG-USP50, but not by the I141R-FLAG-USP50 mutant (Figure 2C), indicating a requirement for USP50 in the restart of HU-stalled forks.

**Figure 2.**
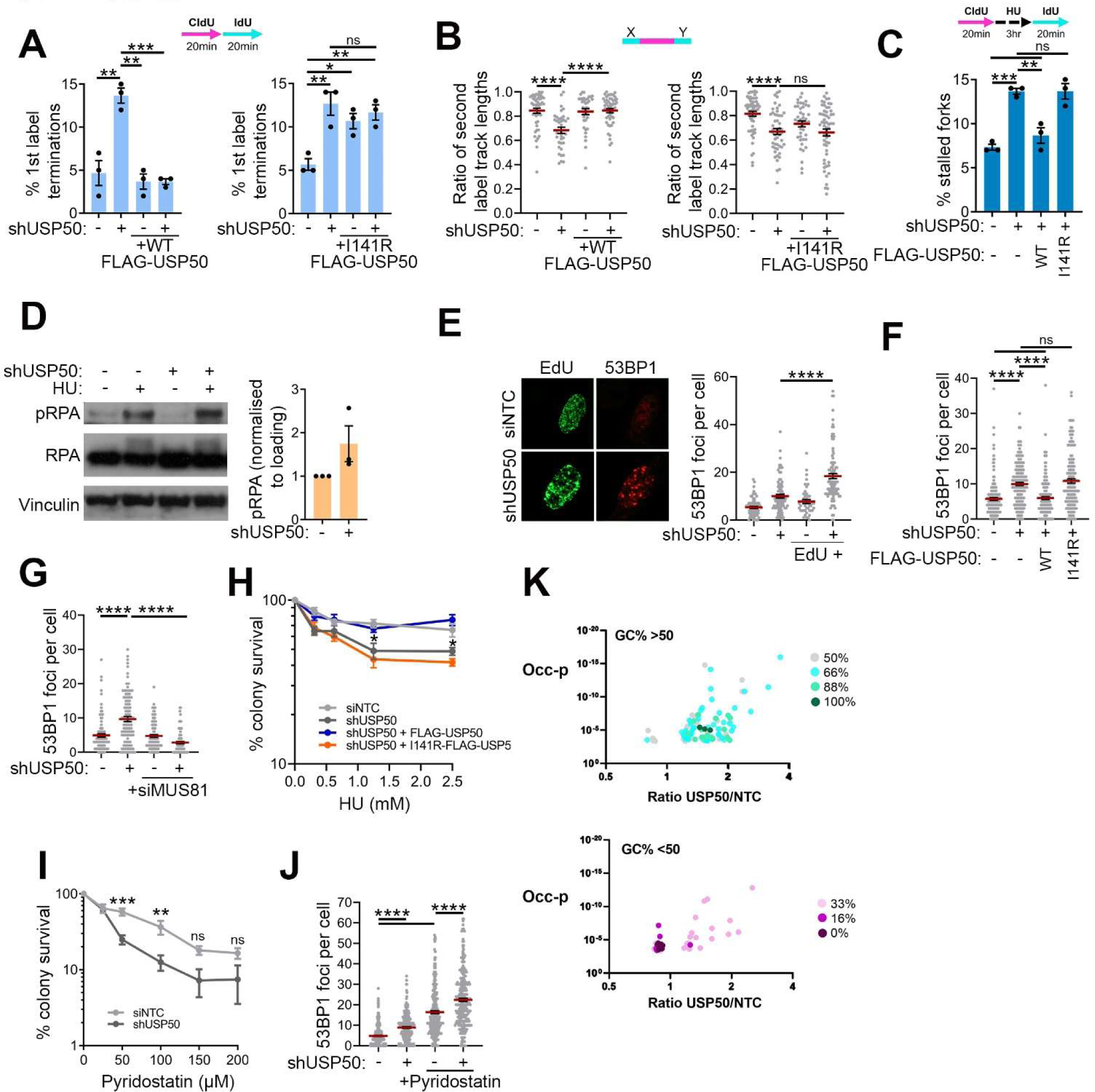
USP50 promotes replication in unperturbed and stressed conditions. A The % of first label terminations from HeLa cells treated with control siRNA (-) or shUSP50 (+) and complemented with FLAG-USP50 (left) or I141R-FLAG-USP50 (right) or uninduced (-). Results are from 3 independent repeats, with n >200 fibres per condition, per repeat. Bars indicate the mean, error bars are SEM. B The IdU tracts within first label origins were measured and the ratio of the left to right lengths were determined as a measure of asymmetry. This was done in HeLa cells treated with control siRNA (-) or shUSP50 (+) and complemented with FLAG-USP50 (left) or I141R-FLAG-USP50 (right) or uninduced (-), quantification is from 3 independent experiments. Red bars indicate the mean, black error bars are SEM. n >35 first label origins measured. C The % of stalled forks from HeLa cells treated with control siRNA (-) or shUSP50 (+) and complemented with FLAG-USP50 or I141R-FLAG-USP50 or uninduced (-). Results are from 3 independent experiments, with n>200 fibres per condition, per repeat. Bars indicate the mean and error bars are SEM. D Western blot (left) and quantification (right) of RPA, pRPA and a vinculin loading control following induction, or not, of shUSP50 in HeLa cells, with and without 3 hours of 5 mM HU treatment. E 53BP1 foci numbers in EdU negative (-) and positive (+) HeLa cells treated with or without shUSP50 and treated with 5 mM HU for 2 hours. Representative images included (left) and quantification (right) is from 2 independent experiments (n=100 cells). Red bars indicate the mean and error bars are SEM. F 53BP1 foci numbers in HeLa cells treated with control siRNA (-) or shUSP50 (+) and complemented with FLAG-USP50 or I141R-FLAG-USP50. Data is from 3 independent experiments (n>150 cells per condition). Red bars indicate the mean and error bars are SEM. G 53BP1 foci numbers in HeLa cells treated with control siRNA (-) or shUSP50 (+) and with control siRNA or siRNA targeting MUS81. Data is from 3 independent experiments (n=100 cells per condition). Red bars indicate the mean and error bars are SEM. H Colony survival following HU treatment (16 hours) was measured in HeLa cells treated with control siRNA (siNTC) or shUSP50 and complemented with FLAG-USP50 or I141R-FLAG-USP50. Data is from 4 independent experiments. Points indicate the mean and error bars are SEM. Statistical analysis done with two-way ANOVA. I Colony survival following Pyridostatin treatment (24 hours) was measured in HeLa cells treated with control siRNA (siNTC) or shUSP50. Data is from 3 independent experiments. Points indicate the mean, error bars are SEM. Statistical analysis done with two-way ANOVA. J 53BP1 foci in HeLa cells treated with control siRNA (-) or shUSP50 (+), with and without treatment with 100 µM Pyridostatin for 24 hours before fixing and staining. Data is from 3 independent experiments (n=250 cells per condition). Red bars indicate the mean and error bars are SEM. K GC-content of 6 bp sequences enriched or reduced (ratio above 1.5 or below 0.8 respectively) in USP50:NTC siRNA treated Hela. Y-axis shows the p-value of the occurrence difference (Occ-p). All significant sequences and p-values are shown in Supplementary Figure 2.

Perturbations in replication fork progression can lead to replicative helicase-polymerase uncoupling, resulting in the accumulation of single-stranded DNA (ssDNA), which is rapidly coated by Replication protein A (RPA), and subsequently phosphorylated ^49^. Upon treatment of cells with USP50 shRNA and HU, we observed a modest increase in pRPA levels (Figure 2D). These findings indicate that USP50 is needed to promote ongoing replication, to aid the protection of stalled forks and to promote replication fork restart.

Prolonged fork stalling can result in the processing of fork structures and the generation of DNA double-strand breaks (DSBs) ^50^. We examined the formation of the DSB marker, 53BP1 foci, in cells pulsed with EdU to allow the labelling of actively replicating cells. We observed increased foci numbers in EdU-positive USP50 shRNA-treated cells and found numbers were further increased upon HU treatment (Figure 2E). Foci were suppressed in USP50 shRNA-expressing cells by the co-expression of FLAG-USP50 but not by the I141R-FLAG-USP50 mutant (Figure 2F). MUS81 is a structure-specific endonuclease subunit that contributes to the cleavage of persistently stalled replication structures ^51, 52, 53^. We found that 53BP1 foci in USP50 shRNA-expressing cells were suppressed by co-depletion of MUS81 (Figure 2G & Supplementary Figure 1E). Together these data correlate poor fork progression and restart in cells deficient for USP50 with increased MUS81-dependent 53BP1 foci, suggesting increased replication fork processing in cells lacking USP50. Depletion of MUS81 did not reduce the fork stalling frequency in cells expressing shUSP50 (Supplementary Figure 1E & F), suggesting that USP50 acts before MUS81 in suppressing fork stalling.

Human *USP50* lies head-to-head with *USP8* on chromosome 15, and the USP50 protein sequence shares 36.6% identity with the C-terminal USP domain of USP8, leading us to consider whether USP8 has a similar function to USP50. Using spontaneous 53BP1 foci to indicate replication difficulties, we compared siRNA sequences targeting USP50 with those targeting USP8. Exposure of cells to siRNA sequences able to deplete USP50, but not those able to deplete USP8, increased 53BP1 foci in otherwise untreated cells (Supplementary Figure 1G-I), suggesting USP8 does not share the ability of USP50 to suppress 53BP1 foci generation.

To address whether USP50 is relevant to the survival of cells experiencing replicative stress, we examined the ability of USP50 shRNA-expressing and complemented cells to form colonies after exposure to HU. USP50 shRNA expression reduced cell survival following HU treatment, which was suppressed by complementation with FLAG-USP50 but not by I141R-FLAG-USP50 (Figure 2H). Further, we observed that USP50 shRNA-expressing cells exposed to the G-quadruplex stabilizing agent pyridostatin also exhibited reduced survival and increased 53BP1 foci formation (Figure 2I & J). These data indicate that USP50 is needed to support the survival of cells undergoing replicative stress.

Our data indicate that USP50 suppresses spontaneous fork collapse. To understand if this occurs at certain genomic regions, we identified the sequences of DSB-proximal sites. To do this, we employed INDUCE-seq. The technique uses adaptors fused to DSB ends, allowing sequencing of 300 to 500 base pairs proximal to the break sites ^54^. We identified 32,448 and 147,395 break sites from 120,000 control and USP50 siRNA-treated cells, respectively, from two technical replicates. The proportion of break-proximal sequences representing short interspersed nuclear elements (SINE), long interspersed nuclear elements (LINE), and long terminal repeat (LTR) elements was lower in USP50 siRNA-treated cells than those treated with control siRNA (Supplementary Figure 2A), suggesting these elements are not the sites sensitive to USP50 loss. We next addressed whether six bp oligonucleotide sequence occurrences at the break-proximal sites differed, finding both significantly enriched and significantly reduced sequences in USP50 siRNA-versus control siRNA-treated cells (Supplementary Figure 2B). Intriguingly, the sequences enriched in USP50 siRNA-treated cells had a GC% of 55.6%, while those reduced had a GC% of 13.3% (Figure 2K). The human genome GC% is 40.85% ^55^, suggesting that USP50 suppresses breakage at some GC-rich regions and contributes to breakage at some AT-rich regions.

### USP50 promotes WRN interactions

To investigate how USP50 influences replication, we first tested a possible role for the Werner RecQ helicase, WRN, which contributes to the replication of GC-rich regions ^56, 57^. Depletion of WRN in shUSP50-treated cells did not further increase the percentage of stalled forks observed after HU treatment (Figure 3A and Supplementary Figure 3A), suggesting that the two proteins function in the same pathway in fork recovery. During replicative stress, WRN interacts with the flap endonuclease FEN1 ^58, 59^, which supports replication after fork stalling ^60^. We noted that the depletion of FEN1 in shUSP50-treated cells similarly did not further increase the percentage of stalled forks observed after HU treatment (Figure 3A and Supplementary Figure 3A). Cells treated with USP50 shRNA showed no change in WRN protein expression levels in whole-cell lysates, but the amount of WRN co-purified with chromatin in HU-treated cells was reduced (Figure 3B). There was no change in FEN1 levels upon USP50 depletion (Supplementary Figure 3B). Following replicative stress, WRN and FEN1 interact with the RAD9/RAD1/HUS1 (9-1-1) checkpoint clamp ^21, 61^. We noted reduced proximity of WRN with both FEN1 and HUS1 of the 9-1-1 complex following USP50 depletion in HU-treated cells (Figure 3C &D). Moreover, we found that the proximity of WRN to FEN1 was improved by complementation with FLAG-USP50 but not by I141R-FLAG-USP50 expression (Figure 3D). These data suggest that both the localisation of WRN to the fork and its subsequent protein-protein interactions are promoted by USP50 and the USP50:Ub interaction.

**Figure 3.**
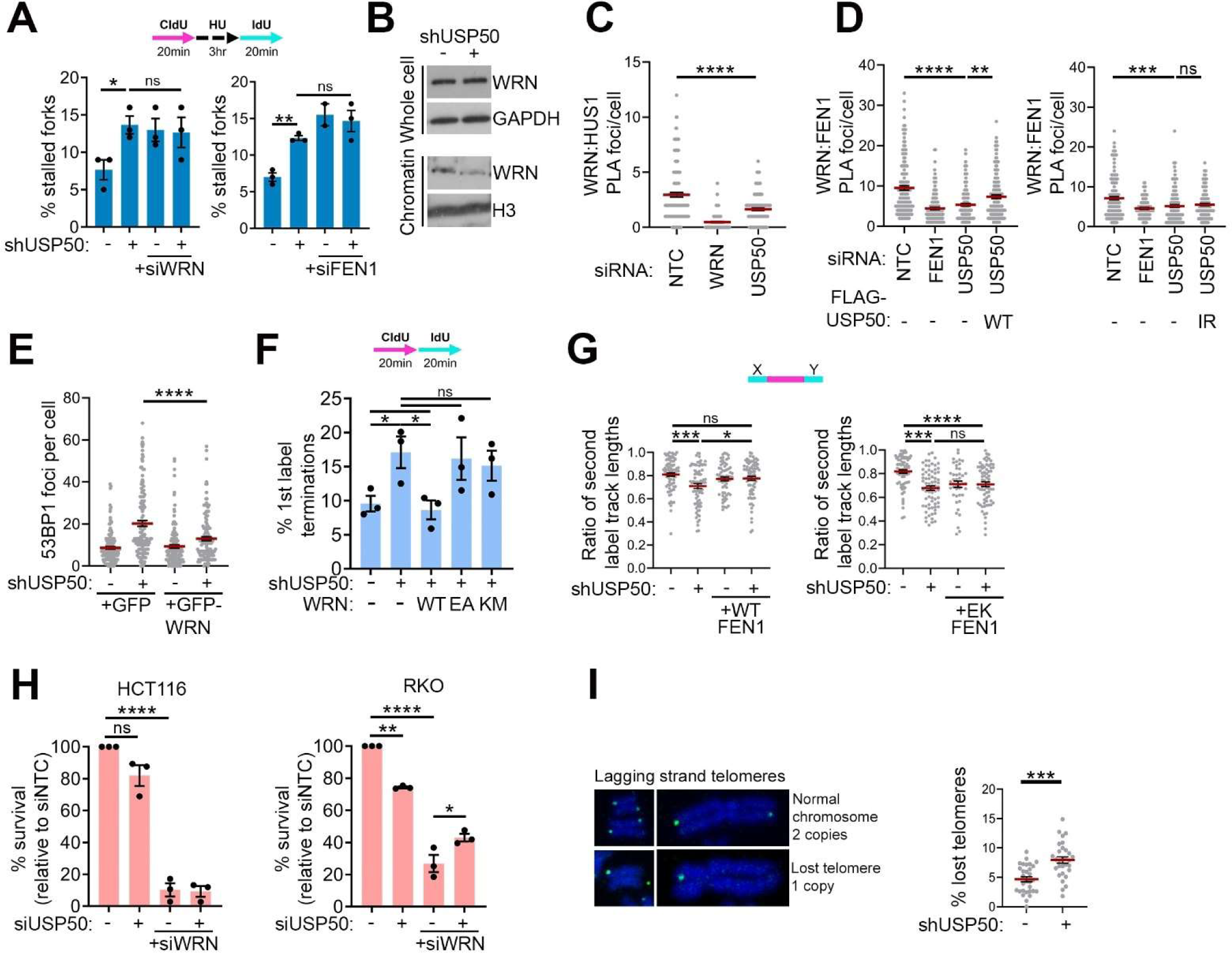
USP50 promotes WRN interactions. A The % of stalled forks from HeLa cells treated with control siRNA (-) or where shUSP50 is induced (+), with treatment with siRNA to WRN (left) or siRNA to FEN1 (right). Results are from 3 independent experiments, with n >200 fibres per condition, per repeat. Bars indicate the mean and error bars are SEM. B Representative blot showing the levels of WRN in the whole-cell lysate and the chromatin-enriched fraction of HeLa cells treated with a control siRNA (-) or shUSP50, followed by 5 mM HU treatment for 3 hours before cell lysis. C WRN proximity to HUS1, part of the 9-1-1 complex, measured via PLA, using antibodies to endogenous WRN and HUS1. HeLa cells treated with control siRNA, siWRN or siUSP50 and then treated with 5 mM HU for 3 hours and scored for red PLA foci. Quantification of PLA foci formation per cell was determined by 3 independent experiments (n >150 cells per condition). Red bars indicate mean and error bars are SEM. D WRN proximity to FEN1 measured via PLA, using antibodies to endogenous WRN and FEN1. HeLa cells treated with control siRNA, siFEN1 or induced shUSP50, complemented with either FLAG-USP50 (left) or I141R-FLAG-USP50 (right). Cells were also treated with 5 mM HU for 3 hours before fixing and scored for red PLA foci. Quantification of PLA foci formation per cell was determined by 3 independent experiments (n >150 cells per condition). Red bars indicate mean and error bars are SEM. E 53BP1 foci in HeLa, EdU positive, cells treated with control siRNA (-) or shUSP50 (+) with expression of GFP or GFP-WRN. Data is from 3 independent experiments (n >130 cells per condition). Red bars indicate the mean and error bars are SEM. F The % of first label terminations from HeLa cells treated with control siRNA (-), or iRNA to USP50, and in cells stably expressing GFP-WT-WRN1, GFP-E84A-WRN or GFP-K577M-WRN. Results are from 3 independent experiments, with n >200 fibres per condition, per repeat. Bars indicate the mean and error bars are SEM. G The IdU tracts within first label origins were measured and the ratio of the left to right lengths were determined as a measure of asymmetry. This was done in HeLa cells treated with control siRNA or shUSP50 and expressing WT Myc-FEN1 (left) or E359K Myc-FEN1 (right), quantification is from 3 independent experiments n>35 first label origins measured. Red bars indicate the mean and error bars are SEM. H Colony survival of HCT116 (left) and RKO (right) cells treated with control siRNA or siUSP50, with and without siRNA to WRN. Data is from 3 independent experiments. Bars indicate the mean and error bars are SEM. Analysed with two-way ANOVA. I Representative CO-FISH image (left), using the c-rich probe for lagging strand replicated telomeres, of metaphase cells treated with telomerase inhibitor and IPTG, or not, to express shUSP50 for 7 days. Graph right shows the percent of telomere loss in lagging strands for cells with and without shUSP50 treatment. Values are mean, bars SEM. n=30 metaphase spreads from 3 independent experiments. Analysed with Welch’s-t-test.

To further test this idea, we over-expressed GFP-WRN and found that 53BP1 foci in shUSP50-treated cells were suppressed (Figure 3E). Consistent with these findings, over-expression of WRN also suppressed the stalling of ongoing forks in USP50 shRNA-treated cells (Figure 3F). Expression of the E84A-WRN mutant that has poor exonuclease function, or the K577M-WRN mutant that perturbs the ATPase/ helicase function of WRN ^62^, failed to suppress ongoing fork stalling (Figure 3F and Supplementary Figure 3C). To probe the WRN:FEN1 interaction further we compared the over-expression of myc-tagged FEN1 (myc-FEN1) with that of the myc-tagged mutant myc-E359K-FEN1. The E359K mutation abolishes the FEN1:WRN interaction and inhibits the gap endonuclease (GEN) activity of FEN1 ^63^. Expression of myc-FEN1 but not the mutant suppressed the occurrence of asymmetric structures from single origins in USP50 siRNA-treated cells, indicating reduced fork stalling following competent myc-FEN1 over-expression (Figure 3G and Supplementary Figure 3D). Independent of WRN, FEN1 has a critical role in Okazaki fragment maturation, and cells without FEN1 activity use poly(ADP-ribose) to recruit XRCC1 in a back-up maturation pathway, which can be observed on PARG inhibition ^64^. We observed no increased poly(ADP-ribose) in cells expressing shUSP50 (Supplementary Figure 3E-H), suggesting Okazaki fragment maturation is unaffected by USP50 loss.

Cancer cells bearing high levels of microsatellite instability (MSI-H) use WRN to replicate expanded (TA)_n_ repeats and are sensitive to loss of WRN, and cancer cells without telomerase expression use WRN and FEN1 to support replication of telomeric repeat (TTAGGG)_6_ of the lagging strand-replicated telomere ^65, 66, 67^. We addressed whether USP50 functions in either of these contexts. We tested MSI-H colon cancer cell lines, HCT116 and RKO, for sensitivity to USP50 siRNA. While these cell lines were susceptible to siRNA targeting of WRN, USP50 siRNA had less impact (Figure 3H). HeLa cells express telomerase, so to examine if USP50 has a role in telomere stability, we grew HeLa cells for seven days in telomerase inhibitor, with or without expression of USP50 shRNA. Metaphase spreads were then subjected to chromosome fluorescence *in situ* hybridization (FISH) using a C-rich probe to detect the G-rich telomere (the strand replicated by lagging strand synthesis). We found that cells with shUSP50 expression showed reduced presence of the G-rich telomeres (Figure 3I). These data suggest that USP50 supports a subset of replicative features associated with WRN function.

### Replication defects in USP50 deficient cells are driven by DNA2 and RECQL4/5

WRN also interacts with the nuclease-helicase DNA2 ^22, 31, 32^, and we anticipated a reduction in DNA2 foci following USP50 depletion as cells recovered from HU treatment. Surprisingly, however, although we observed no change in total DNA2 protein (Supplementary Figure 3I), we observed increased DNA2 foci (Figure 4A). To test whether this finding relates to the defects in fork kinetics observed in shUSP50-treated cells, we co-depleted USP50 and DNA2. Contrary to expectations, we found that the depletion of DNA2 improved fork restart, suppressed ongoing fork stalling, and reduced fork asymmetry in cells treated with shUSP50 (Figure 4B-D and Supplementary Figure 3I). DNA2 depletion also restored fork restart in cells overexpressing the I141R-FLAG-USP50 mutant (Figure 4E). Furthermore, DNA2 siRNA treatment also suppressed the appearance of increased pRPA in shUSP50-treated cells (Figure 4F). We utilized the selective DNA2 nuclease inhibitor C5, which inhibits DNA binding and nuclease function ^68^, and examined the number of restarted forks. 20 μM C5 treatment suppressed the restart defect of shUSP50-treated cells (Figure 4G). In contrast, inhibition of the nuclease MRE11, present at forks and also implicated in restart ^69^, had no impact (Supplementary Figure 3J). These data suggest that DNA2 nuclease activity is responsible for the replication defects observed in cells deficient for USP50.

**Figure 4.**
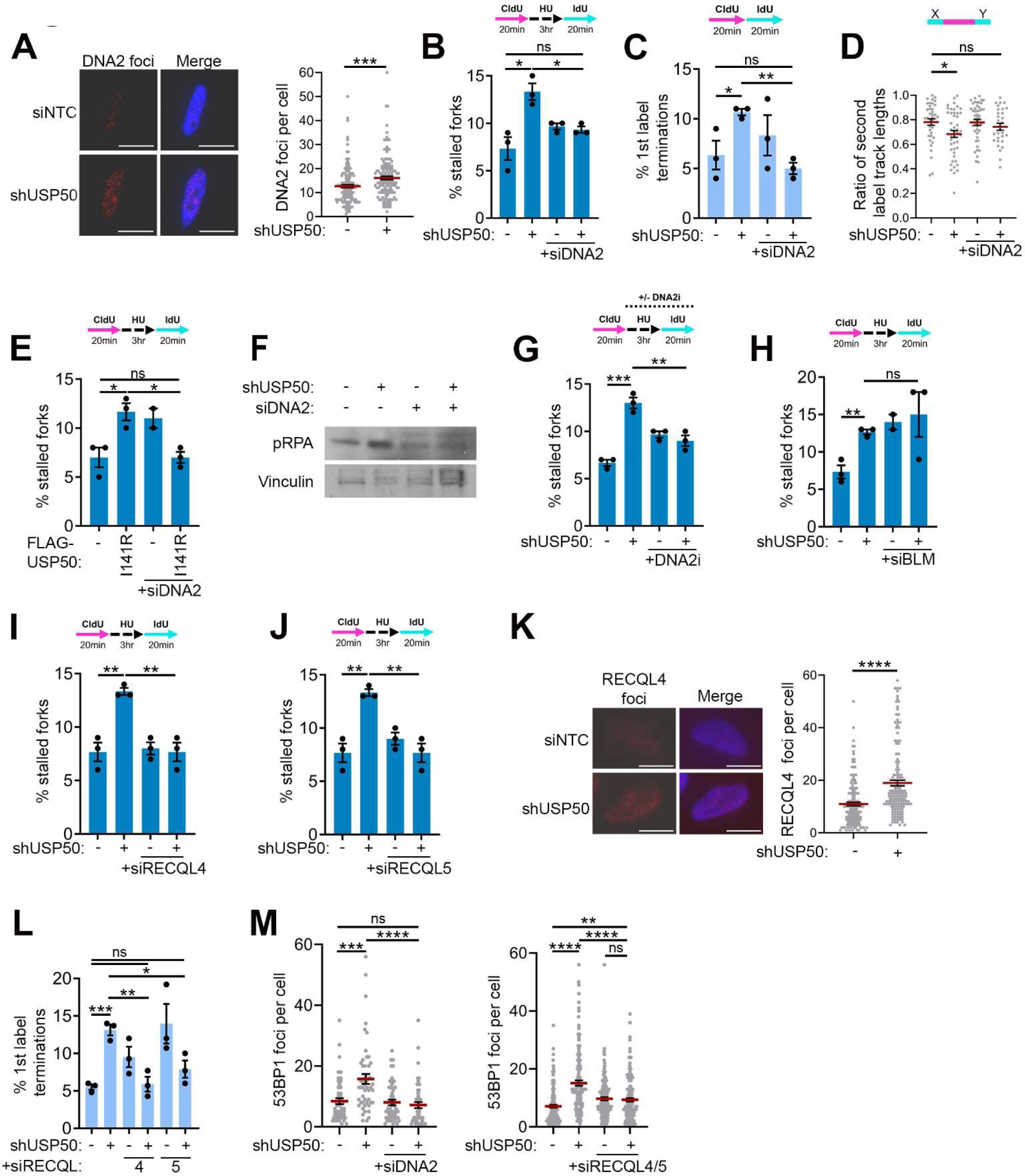
Replication defects in USP50 deficient cells are driven by DNA2 and RECQL4/5. A DNA2 foci numbers in HeLa, EdU positive, cells treated with control siRNA (-) or shUSP50. Cells were treated with 5 mM HU for 3 hours, washed once and allowed to recover for 30 min before fixation. Representative images (left, scale bar is 10µm) are shown along with data (right) from 3 independent experiments (n >150 cells per condition). Red bars indicate the mean and error bars are SEM. B The % of stalled forks from HeLa cells treated with shUSP50, with and without co-depletion of DNA2. Data is from 3 independent experiments, with n>200 fibres per condition, per repeat. Bars indicate the mean and error bars are SEM. C The % of first label terminations from HeLa cells treated with shUSP50, with or without co-depletion of DNA2. Results are from 3 independent experiments, with n>200 fibres per condition, per repeat. Bars indicate the mean and error bars are SEM. D The IdU tract lengths for first label origins were measured and the ratio of second label to first label lengths were determined as a measure of asymmetry. This was done in HeLa cells treated with control siRNA or shUSP50 with and without co-depletion of DNA2. Quantification is from 3 independent experiments (n>35 per condition). Red bars indicate the mean and error bars are SEM. E The % of stalled forks from HeLa cells treated with control siRNA or shUSP50, with and without complementation of I141R-FLAG-USP50, with and without depletion of DNA2. Results are from 3 independent experiments, with n>200 fibres per condition, per repeat. Bars indicate the mean and error bars are SEM. F Western blot showing the levels of pRPA (S4/8) following treatment with a control siRNA (-) or shUSP50 (+), with and without co-depletion of DNA2. Cells were treated with 5 mM HU for 3 hours prior to being lysed. Vinculin is also shown as a loading control. G The % of stalled forks from HeLa cells treated with control siRNA (-) or shUSP50 (+), with and without co-treatment of a C5 DNA2i (20 µM). Results are from 3 independent repeats, with n>200 fibres per condition, per repeat. Bars indicate the mean and error bars are SEM. H The % of stalled forks from HeLa cells treated with shUSP50, with and without co-depletion of BLM. Results are from 3 independent repeats, with n>200 fibres per condition, per repeat. Bars indicate the mean, error bars are SEM. I The % of stalled forks from HeLa cells treated with control siRNA or shUSP50, with and without co-depletion of RECQL4. Results are from 3 independent repeats, with n>200 fibres per condition, per repeat. Bars indicate the mean and error bars are SEM. J The % of stalled forks from HeLa cells treated with control siRNA or shUSP50, with and without co-depletion of RECQL5. Results are from 3 independent repeats, with n>200 fibres per condition, per repeat. Bars indicate the mean and error bars are SEM. K RECQL4 foci numbers in HeLa, EdU positive, cells treated with control siRNA (-) or shUSP50. Cells were treated with 5 mM HU for 3 hours, washed once and allowed to recover for 30 min before fixation. Representative images (left, scale bar is 10µm) are shown along with data (right) from 3 independent experiments (n >150 cells per condition). Bars indicate the mean and error bars are SEM. L The % of first label terminations from HeLa cells treated with shUSP50, with or without co-depletion of RECQL4 or RECQL5. Results are from 3 independent experiments, with n>200 fibres per condition, per repeat. Bars indicate the mean and error bars are SEM. M 53BP1 foci numbers in HeLa cells treated with control siRNA (-) or shUSP50 (+) and with siRNA targeting DNA2 (left) or RECQL4 and 5 (right). Cells were pulsed with EdU, treated with 5mM HU for 3 hrs and given a 20-minute recovery period without HU before fixation. Data is from 2 independent experiments (n>58 cells per condition). Bars indicate the mean and error bars are SEM.

To process dsDNA, DNA2 requires a companion helicase such as WRN or BLM^31, 32, 70^. Considering the surprising finding that DNA2 contributes to replication defects in USP50-depleted cells, where WRN association with forks is reduced, we next assessed whether an alternative supporting helicase contributes to replication defects. Depletion of BLM slightly increased the proportion of stalled replication structures after release from HU in the presence of USP50 shRNA (Figure 4H and Supplementary Figure 3K), suggesting that BLM is not responsible for poor fork recovery of shUSP50-treated cells. Similarly, and as expected ^15, 16, 22^, siRNA to RECQL1 also further suppressed fork restart (Supplementary Figure 3K & L). These data suggest that BLM and RECQL1 work in pathways separate from USP50 in fork restart. In contrast, co-depletion of either RECQL4 or RECQL5 with shUSP50 treatment reduced the number of stalled forks after release from HU treatment (Figure 4I, 4J and Supplementary Figure 3K), suggesting that these proteins have a similar role to DNA2 in suppressing fork recovery in the context of USP50 loss. Consistent with these findings, we noted that RECQL4 showed increased foci formation in USP50 shRNA -treated cells recovering from HU exposure (Figure 4K). When we examined ongoing forks, we similarly noted that co-depletion of either RECQL4 or RECQL5 suppressed on-going fork stalling of cells treated with USP50 siRNA (Figure 4L), suggesting these proteins also contribute to spontaneous stalling of forks. Intriguingly RECQL5 depletion, and to a lesser degree RECQL4 depletion, alone increased fork stalling, and since stalling was reduced when co-depleted with USP50, these data suggest a deleterious role for USP50 when RECQL4 or RECQL5 are absent. Finally, we assessed the impact of the RECQL helicases and DNA2 on spontaneous 53BP1 foci as a measure of collapsed replication forks. Intriguingly, 53BP1 foci were not increased following depletion of the helicases or DNA2 alone, whereas the elevated 53BP1 foci observed on shUSP50 treatment were suppressed by co-depletion of RECQL4 and RECQL5 or DNA2 (Figure 4M).

These data suggest that the DNA2 nuclease and RECQL4/5 helicases contribute to replication defects observed in cells deficient for USP50.

## Discussion

USP50 has been implicated in inflammasome signalling, erythropoiesis, the G2/M checkpoint, and Human Growth-Factor-dependent cell scattering ^71, 72, 73, 74^. Mice homozygous for disrupted *Usp50* (*Usp50^tm1(KOMP)Vlcg^*) die *in utero* ^75^, indicating it is required for life, and two previous siRNA screens have identified USP50 as a candidate replication-related protein ^41, 42^. Here we reveal the surprising finding that this lowly-expressed Ub-binding protein suppresses alternative RecQ helicase use and deleterious DNA2 activity during replication. Depletion of USP50 reduces WRN:FEN1 presence at stalled forks, increases resection measures and DNA2 and RECQL4 foci as forks recover, and results in the suppression of fork restart by DNA2 and RECQL4/5. Ongoing replication also requires USP50 to suppress the harmful impact of DNA2, RECQL4 and RECQL5. As 53BP1 foci occurrence, and by inference fork collapse, correlates more with the effects of the helicases on fork restart than on the proportion of stalled ongoing forks, we suggest it is the impact of USP50 loss and DNA2:RECQL4/5 activity on suppressing fork restart that drives fork collapse.

*In vitro* WRN can unwind various structures, some of which are likely to occur ahead of the replication fork and others at or behind the fork junction. The WRN helicase unwinds the chicken-foot intermediate associated with regressed replication forks and 3′-tailed duplexes, bubble structures, forked duplexes, G-quadruplex structures, and DNA displacement loops ^76, 77^. Intriguingly we find that USP50 function aligns with some functions of WRN but not others. Like WRN^18^, USP50 supports ongoing replication and suppresses MUS81-mediated fork collapse. Further, like WRN:FEN1^63, 65, 66^, it promotes the stability of lagging, G-rich telomeres. We find that USP50 suppresses DBSs near some GC-rich sequences, but the precise type of DNA structure that USP50 supports is not yet clear. For example, while more break sites in USP50-depleted cells overlap with G-quadruplex mapped sequences^78, 79, 80^ than control siRNA-treated cells (2648 *Vs* 1726), these do not represent an increased proportion of the breaks (1.79% *Vs* 5.3% respectively), indicating that G-quadruplex forming sequences are not over-represented near the breaks found in USP50 depleted cells. Moreover, USP50 depletion is not lethal to MSI-H cells, suggesting that USP50 does not relate to the ability of the WRN helicase to process the non-B form (TA)_n_ DNA repeats ahead of the replication fork that are thought to be the basis of the WRN:MSI-H synthetic lethality ^13^. In contrast, the depletion of USP50 suppressed the appearance of DSBs near AT-rich DNA sequences in microsatellite-stable HeLa cells. Thus the data support the view that USP50 supports a subset of WRN functions.

Our data reveal a differential influence of USP50 on WRN partner nucleases FEN1 and DNA2, in which USP50 supports FEN1 and suppresses DNA2 activity. We find WRN interaction with FEN1 is promoted by USP50, and both WRN and FEN1 over-expression minimize the impact of USP50 loss. Increased expression of the FEN1 nuclease may increase WRN stability at stalled forks, and/or provide FEN1 GEN activity as suggested by the inability of the E359K-FEN1 mutant to restore symmetrical forks. Like WRN loss, FEN1 depletion is epistatic with shUSP50 treatment in fork restart, consistent with the promotion of WRN-FEN1 activities by USP50. Also consistent with the role of USP50 in promoting WRN:FEN1 interactions, both WRN:FEN1 interaction and GEN activity are required for lagging strand telomere stability ^66^.

In contrast to the impact on FEN1, loss of USP50 results in increased DNA2 recruitment and we find that DNA2, unrestrained by USP50, can suppress ongoing replication, cause fork asymmetry, suppress fork restart and promote fork collapse. In normal replication, the promotion of regressed replication fork restart involves DNA2:WRN^22^, yet in the absence of USP50, WRN and DNA2 loss are no longer epistatic in fork restart. The interaction faces of FEN1 and DNA2 have been mapped to large and overlapping regions of WRN^32, 59^, but whether FEN1:WRN and DNA2:WRN interactions and roles are sequential, co-dependent or differ depending on the stalled structure is not currently known. As increased DNA2 presence and activity and decreased WRN is observed at stalled forks after USP50 depletion, it would appear that DNA2 does not necessarily require the function of WRN in this context.

Remarkably, in the absence of USP50, RECQL4/5 helicases contribute to the stalling of ongoing forks, suppress fork restart, and promote replication fork collapse. We observed increased RECQL4 foci in USP50 depleted cells recovering from HU-mediated fork arrest, and it is perhaps relevant that RECQL5 is retained at damage sites longer in cells derived from Werner’s syndrome patients^24^. Thus, it is possible that the reduced WRN localisation to stalled forks, observed following USP50 depletion, contributes to the increased RECQL5/4 use. We suggest the simplest model for the role of DNA2 and RECQL4/5 helicases in replication fork problems when USP50 is depleted reflects an altered balance from WRN towards RECQL4/5 helicase involvement correlating with increased DNA2-mediated resection, slowed restart and increased MUS81-mediated fork collapse (illustrated in Figure 5). We recognise there are several potential alternative models, from inappropriate strand annealing mediated by RECQL4/5, (which show relatively weak helicase activity and relatively strong strand annealing ^31^ ^81, 82^), to a failure to destabilise ssDNA secondary structures (which WRN or BLM, but not other RecQ helicases, can achieve ^83^). RECQL4 and RECQL5 function in replication initiation ^84, 85^, homologous recombinations and the resolution of replication: transcription conflicts ^86, 87^, respectively. Previous reports have suggested RECQL5 can support replication in specific circumstances^23,24^. Our observations indicate that RECQL5 and RECQL4 can disrupt replication restart and ongoing replication if not adequately controlled.

**Figure 5.**
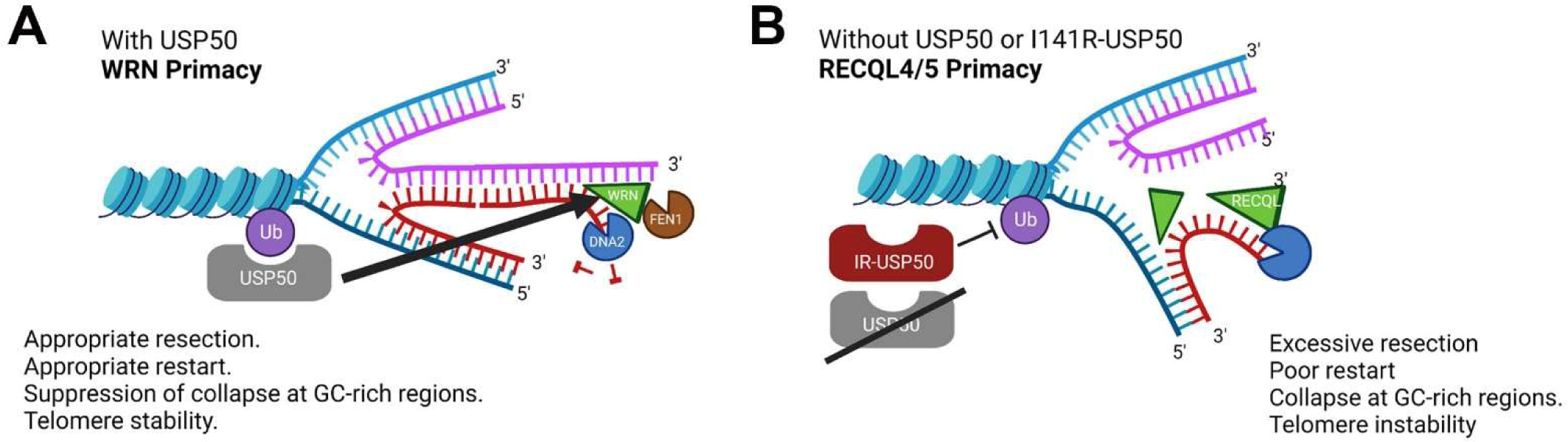
USP50 regulation of WRN and RECQL4/5 helicases. A USP50 is recruited to stalled forks through a Ub-interaction and supports a subset of WRN functions at restarting forks, including FEN1-associated functions. B In USP50 deficient cells or cells expressing I141R-USP50, WRN recruitment is reduced and inappropriate RECQL4/5 function and DNA2 activity occurs, resulting in excessive resection. This loss of USP50 leads to suppression of fork restart, telomere instability and fork collapse. (Created with BioRender.com)

Approximately 10% of known mammalian deubiquitinating enzymes are predicted to be inactive, ‘pseudo-enzymes’. The USP-class of pseudo-enzymes with known cellular activities have functions attributed to domains other than their USPs^88^. While active site oxidation can inhibit enzymatic activity without preventing Ub-binding and active USP enzymes can be converted into Ub-binding proteins experimentally by active site substitutions ^89^ ^90^, the current study is the first, to our knowledge, demonstrating that the Ub-binding of a catalytically inactive DUB is critical to its function.

By revealing a critical role for USP50 in repressing DNA2 activity and promoting canonical RecQ-helicase use, the current data suggests new questions, the investigation of which will inform how mammalian DNA replication is facilitated. These questions include: Whether USP50 regulates a process able to amplify its effect (such as a kinase or an E3 ligase) or acts stoichiometrically? Does the Ub modification of a specific substrate or wide-spread chromatin modification at stalled fork structures ^91, 92, 93^ drive USP50 localisation to stalled forks? How does USP50 promote WRN localisation to stalled replication structures, and does suppression of DNA2 relate to WRN changes or occur independently? Finally, our data predict a further as-yet-unknown component since over-expression of USP50 lacking the ability to bind Ub can suppress normal replication kinetics without co-treatment with USP50 shRNA. What is this component?

In summary, we identify a new Ub-mediated pathway, centred on USP50, that influences WRN, FEN1, DNA2, RECQL4 and RECQL5 at ongoing and stalled replication forks. Our data reveal that the canonical use of WRN and its partners, the suppression of the RECQL4 and RECQL5 helicases and the prevention of aberrant DNA2 activity requires USP50. Further investigation of this pathway will offer opportunities to understand better these proteins’ roles in replication and human disorders.

---

**Supplementary Figure 1.**
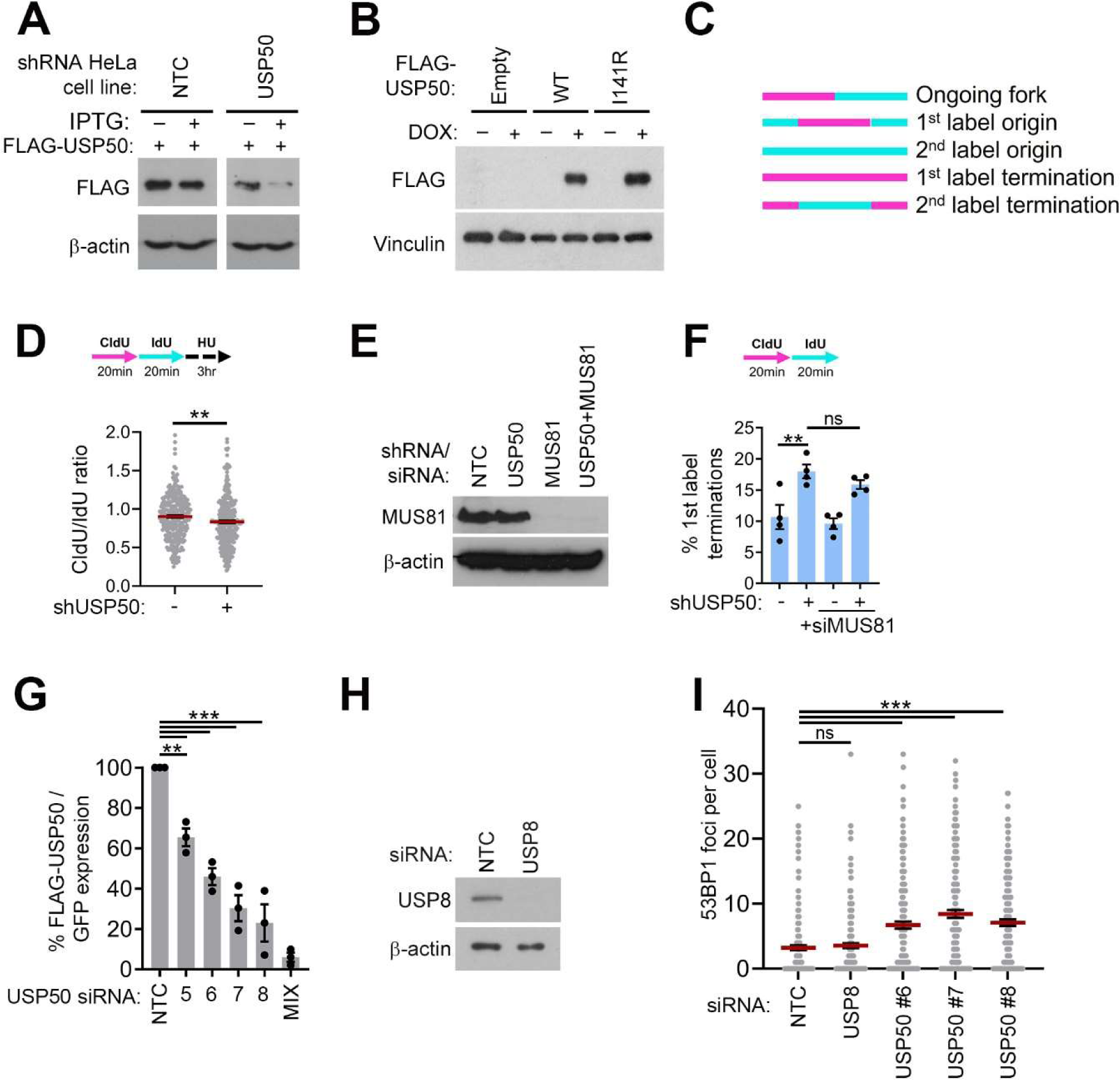
A FLAG western blot of two Hela cell lines expressing wild-type FLAG-USP50, the first parental line was treated with siRNA to Luciferase as a non-targeting control (siNTC), with and without isopropyl-β-D-1-thiogalactopyranoside (IPTG) and the second was a line generated to express an IPTG-inducible shRNA to USP50, treated with and without IPTG. B FLAG western blot showing of Hela cell lysates bearing inducible shUSP50 and of lysates from cell lines also bearing doxycycline-inducible, shRNA-resistant FLAG-USP50 wild-type at the protein level (WT) or additionally bearing I141R, treated of not with doxycycline (Dox). All cells were also treated with IPTG. C Diagram to illustrate structures identified in the DNA fibre assay to measure fork kinetics by assessing CldU and IdU labelling. D The CldU and IdU tract lengths for ongoing forks were measured and the ratios were determined as a measure of for protection. This was done in HeLa cells treated with control siRNA or shUSP50. Quantification is from 3 independent experiments (100 fibres per condition, per repeat). Red bars indicate the mean and error bars are SEM. E Western blot to verify the endogenous depletion of MUS81 by siRNA treatment, with and without treatment of shUSP50. F The % 1^st^ label terminations were measured in HeLa cells treated with siRNA control or shUSP50, with and without treatment with MUS81 siRNA. Results are from 4 independent experiments, n>200 fibres per condition per repeat. Bars indicate the mean and error bars are SEM. G Measurement of non-shRNA (or siRNA) resistant FLAG-USP50 relative to co-expressed GFP expressed as a % of FLAG-USP50 in cells treated with non-targeting control siRNA (NTC), or with various USP50 siRNA sequences. Bars indicate the mean and error bars are SEM. **p<0.01, ***p<0.001, two-tailed ANOVA. H Western blot assessment of USP8 expression in cells treated with non-targeting control siRNA (NTC) or siRNA to USP8. I 53BP1 foci numbers in HeLa cells treated with control siRNA, siRNA targeting USP8, or the numbered USP50 siRNAs. Results from 2 independent experiments (n=200 cells per condition). Bars indicate the mean and error bars are SEM. ***p<0.001 and ns = non-significant Kruskal-Wallis 1-tailed test

**Supplementary Figure 2.**
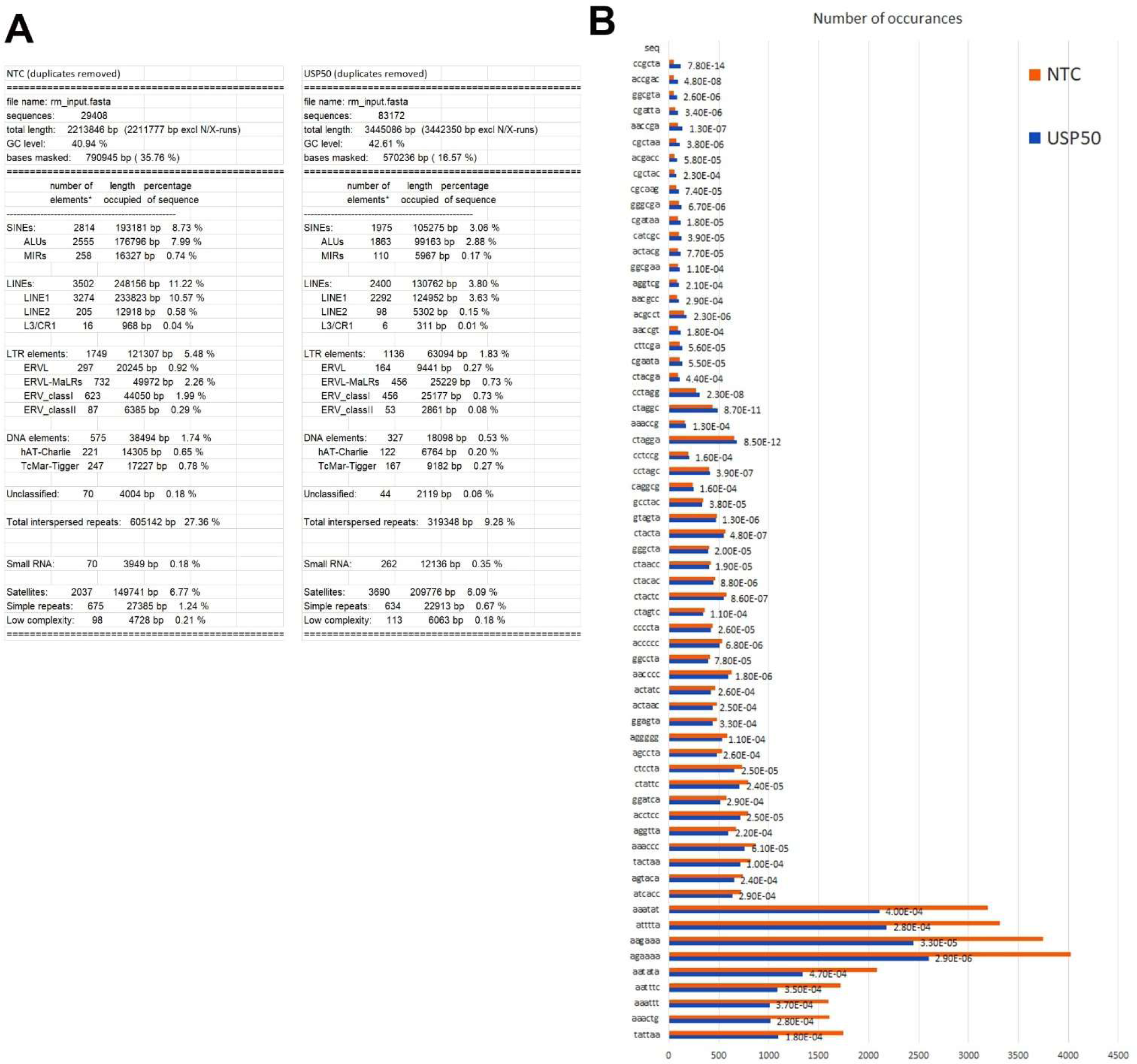
A RepeatMasker statistics on sequences proximal to breaks in cells treated with non-targeting control siRNA (NTC) or siRNA targeting USP50. B Frequency of significantly different occurrences of sequences (6 bp) near a double strand break site between shUSP50 (blue) and siNTC (orange) treated HeLa cells. Sequences are shown on the y-axis and the number of occurrences on the x-axis. Numbers on right of bars show p value.

**Supplementary Figure 3.**
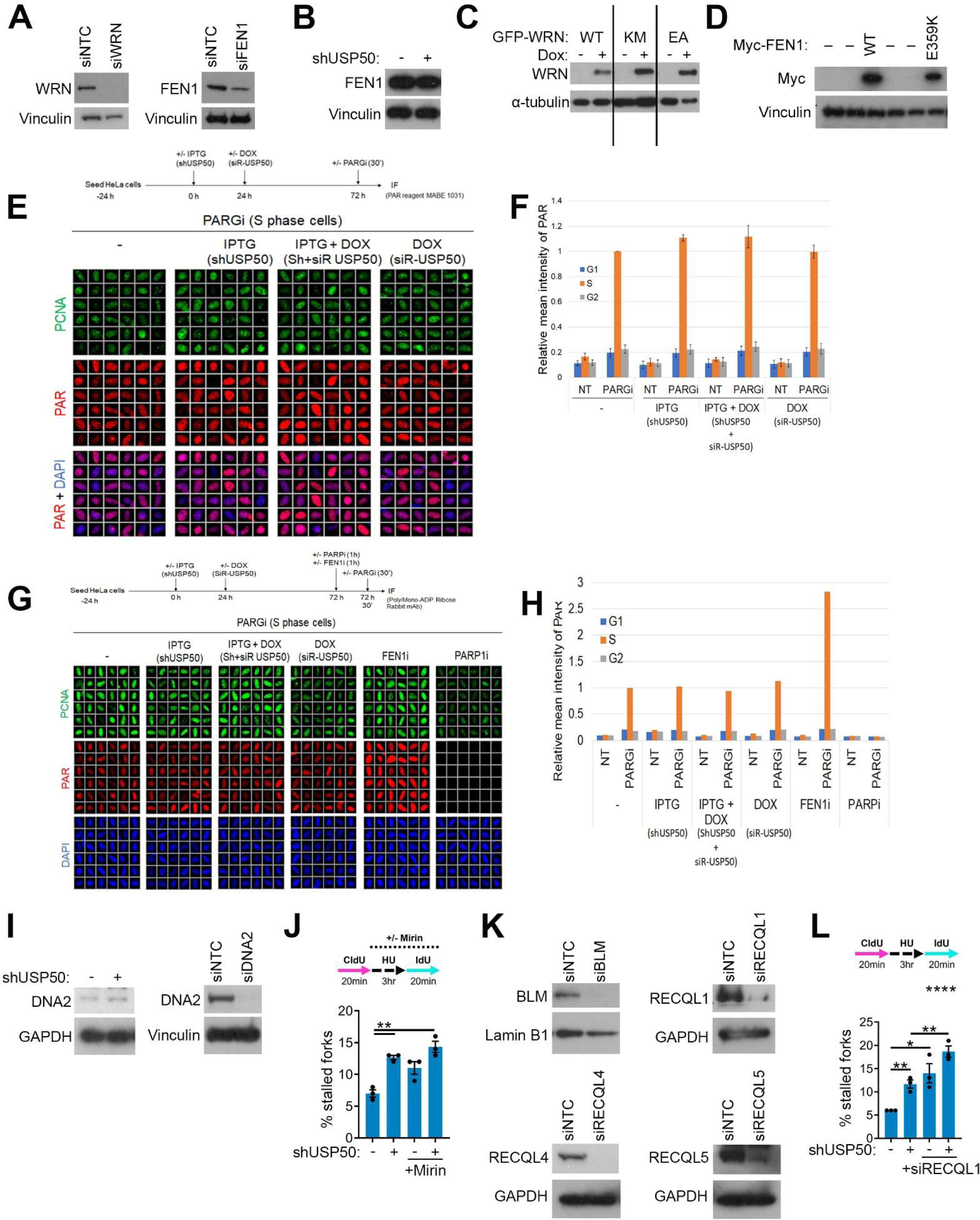
A Western blot analysis of lysate from cells treated with non-targeting control siRNA (luciferase) siNTC or siRNA to WRN (left) or FEN1 (right). Blots were probed for WRN and FEN1 respectively and both were also probed for Vinculin as a loading control. B Western blot analysis of lysate from cells treated with non-targeting control siRNA (luciferase) siNTC or shUSP50. Blots were probed for FEN1 and Vinculin as a loading control. C Western blot analysis of cells bearing genomically integrated GFP-WRN variants, WT-WRN, E84A-WRN (EA) or K577M-WRN (KM), induced by doxycycline. Lysates were blotted and incubated with antibodies to loading control α-tubulin and to WRN. D Western blot analysis of cells bearing genomically integrated, siRNA resistant myc-FEN variants; WT-FEN1, and E395K-FEN1. Lysates were blotted and incubated with antibodies to loading control vinculin and to myc. E (Top) A schematic illustration of the experimental setup for the detection of PAR using the poly-ADP-ribose binding reagent MABE1031. (Bottom) Representative ScanR images. F Quantification of mean fluorescence intensities of PAR in HeLa cells incubated with IPTG, Doxycycline (DOX) or both from 3 independent experiments. 10 µM PARG inhibitor was added or not during the last 30 min of incubation with the mentioned above compounds. S phase cells were identified based on PCNA staining. G1 and G2 phase cells were distinguished based on DAPI intensity. The mean fluorescent intensity of PAR for each sample was normalised to the mean fluorescent intensity of S phase PAR in the presence of PARGi, but without any other treatments. Bars indicate the mean and error bars are SEM. G (Top) Schematic illustration of the experimental setup for the detection of PAR.(Bottom) Representative ScanR images. H Quantification of mean fluorescence intensities of PAR in HeLa cells incubated with IPTG, DOX or both. Untreated cells were incubated with 10 µM FEN1i (positive control) or 10 µM PARPi (negative control). 10 µM PARGi was added or not during the last 30 min of incubation with the mentioned above compounds. I Western blot analysis of lysate from cells treated with non-targeting control siRNA (luciferase) siNTC and either shUSP50 (left) or siDNA2 (right). Blots were probed for DNA2 and either GAPDH or Vinculin as a loading control. J The % of stalled forks from HeLa cells treated with non-targeting siRNA or shUSP50, with and without co-treatment of 50 µM Mirin. Data is from 3 independent experiments, with n>200 fibres per condition per repeat. Bars indicate the mean and error bars are SEM. K Western blot analysis of lysates from cells treated with non-targeting control siRNA (siNTC) or siRNA to a family member of the RecQ helicases. Included are siBLM (top left), siRECQL1 (top right), siRECQL4 (bottom left) and RECQL5 (bottom right). Blots were probed for the indicated RecQ helicase family member and Lamin B1 or GAPDH as loading controls. L The % of stalled forks from HeLa cells treated with non-targeting control siRNA or shUSP50, with and without co-depletion of RECQL1. Data is from 3 independent experiments, with n>200 fibres per condition per repeat. Bars indicate the mean and error bars are SEM.

**Supplementary Figure 4.**
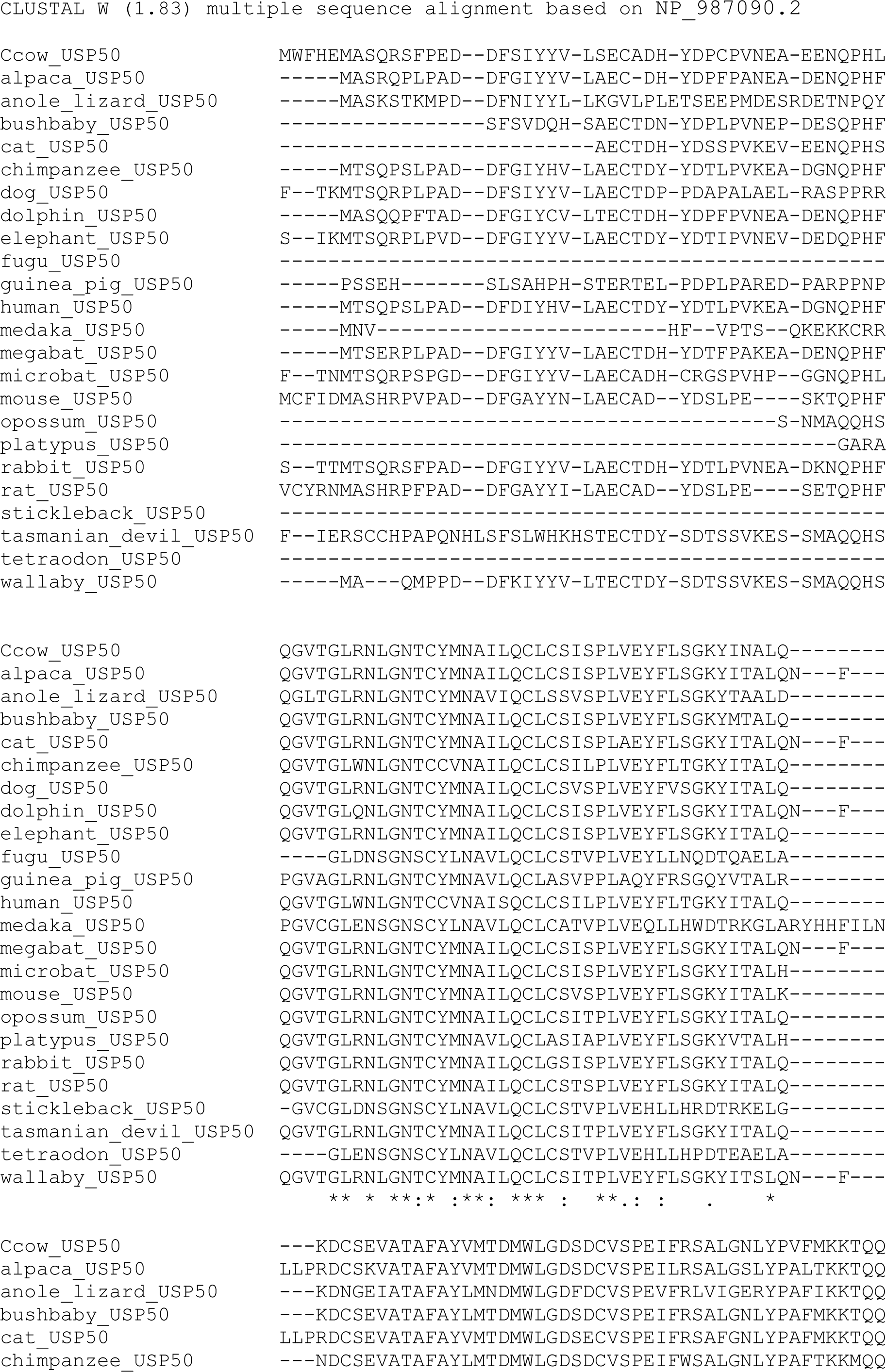
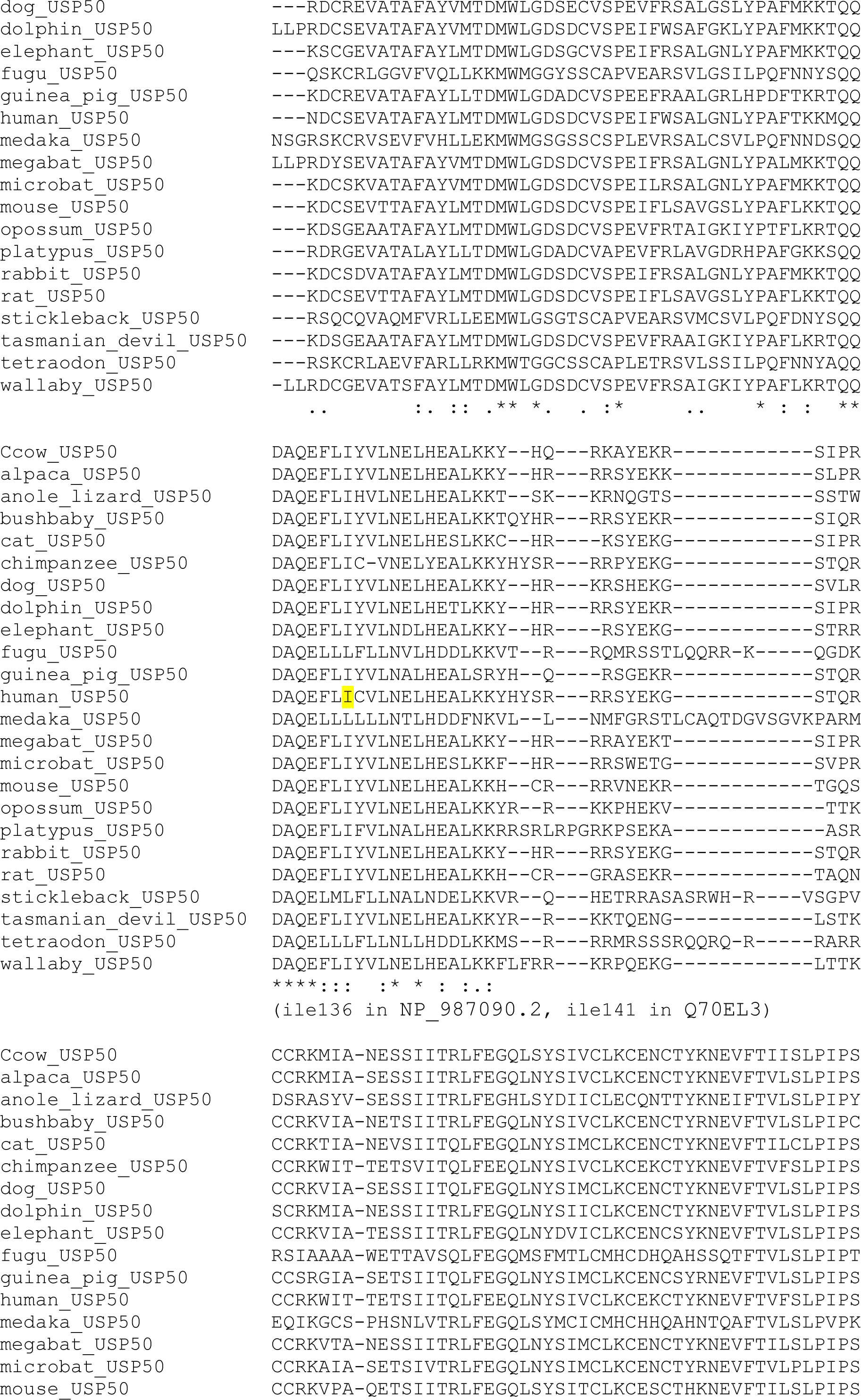
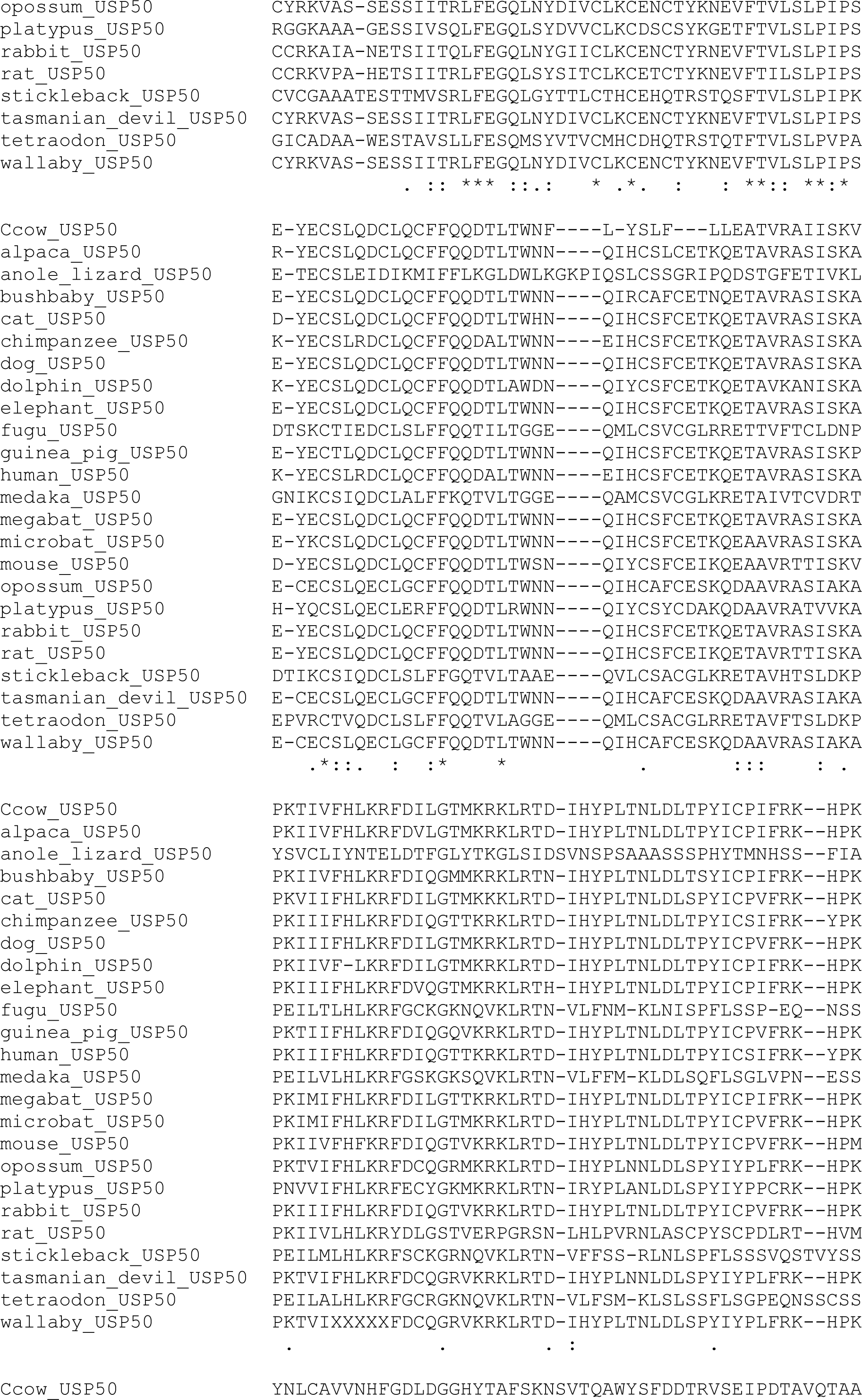
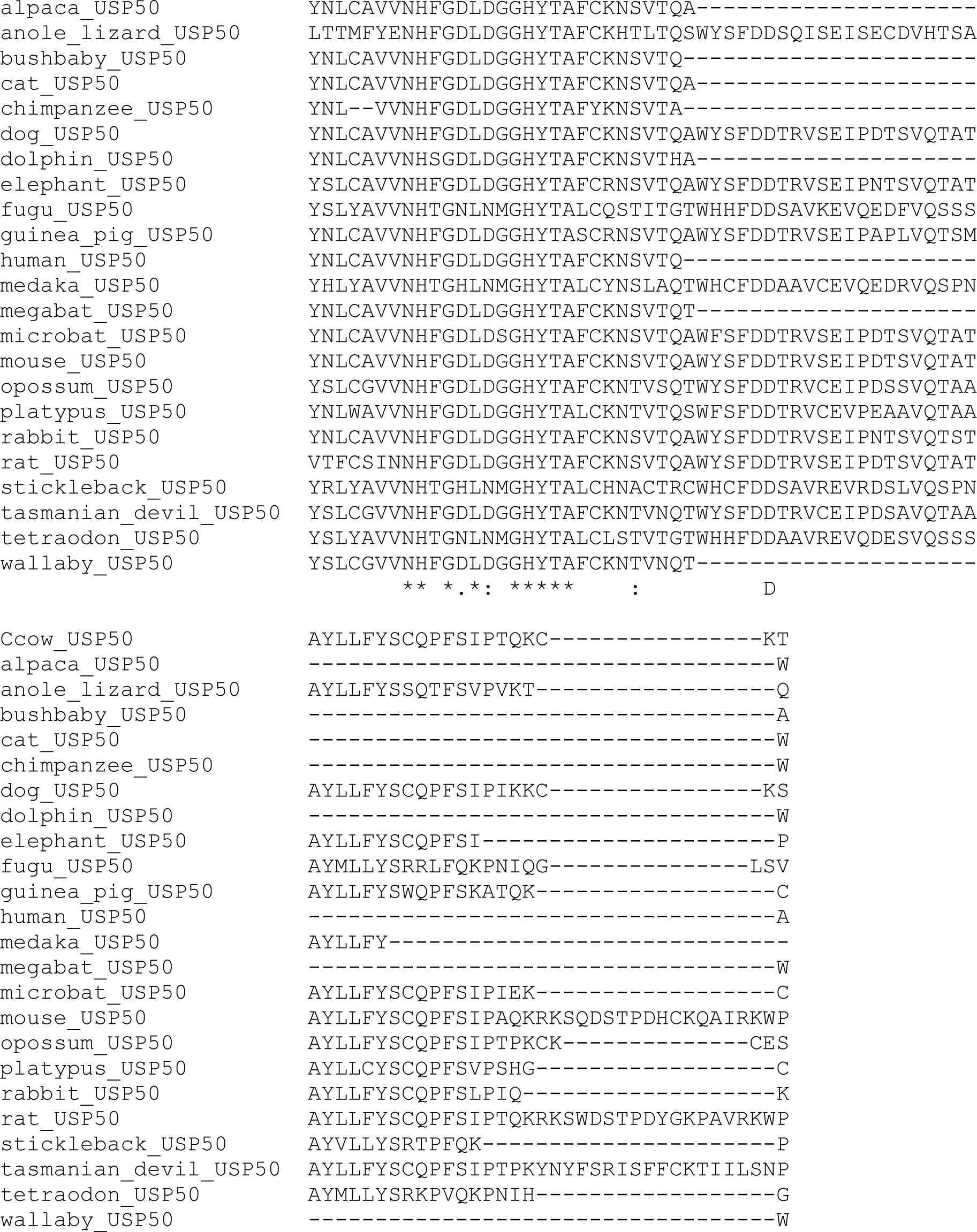

## Acknowledgments

Grant funding was as follows. CRUK: C8820/A28283 and A28283 (HLM, HRS, GR, KS, and JB), Breast Cancer Now: 2015MayPR499 (KS) and Studentship 2010NovPhD02 (HRS). Wellcome Trust 206343/Z/17/Z (AW and AG), CRUK Centre training (KE) and University of Birmingham (SV). Worldwide Cancer Research 13-1048 (EP) and Medical Research Council MR/J007595/1 (EP). FLAG-HA-USP50 was a gift from Wade Harper (Addgene plasmid # 22588; http://n2t.net/addgene:22588; RRID:Addgene22588) ^94^. Nick Davies provided expert advice for the COFISH experiments. We thank Professor Andrew Beggs, University of Birmingham for HCT116 and RKO cells, and Professor Keith Caldecott for discussions and reading the manuscript. We thank the Microscopy and Imaging Services at Birmingham University (MISBU) and the UoB Flow Cytometry Services (UoBFC) in the Tech Hub facility for microscope and FACS support and maintenance.

## Author Contributions

HRS generated reagents, fibre experiments and *in vitro* analysis. HM performed fibre assays, chromatin fractionations, immunoprecipitations, and PLA analysis. GR performed Chromatin fractionation assays. KS and AW performed PLA analysis. SV generated reagents and performed foci experiments. KE performed fibre experiments and colony assays. AJG generated reagents, performed cell survival and foci analysis. JB provided technical support. AK performed the PARGi experiments. EP and EJB advised HRS. FD and PVE performed INDUCE-Seq. SR supervised FD and PVE. MS advised on bioinformatics. RMD performed CO-FISH analysis. JRM, HM and RMD wrote the paper. All authors commented on the paper and research. JRM conceived and directed the project.

## Materials and Methods

### Tissue culture

HCT116 were grown in McCoy’s 5A medium with 10% Fetal Bovine Serum (FBS) + 1% Penicillin/Streptomycin (PS). RKO cells were grown in Minimum Essential Medium with 10% FBS, 2 mM L-glutamine and 1% PS. These cell lines were sourced from Professor Andrew Beggs, University of Birmingham. HeLa cells were grown in Dulbecco’s Modified Eagle Media supplemented with 10% FBS and 1% PS. Cells were cultured in Corning T75 flasks and 10 cm2 plates and kept at 5% CO2 and 37°C. Details of key chemicals are in **Supplementary Table 3**.

### Inducible shRNA

Custom Lentiviral shRNA USP50 sequence (based on the USP50-7 siRNA sequence C UAC CCA GCA UUU ACG) or non-targeting control sequence cloned into the pLKO-puro-IPTG-3xLacO vector were made by Sigma-Aldrich (Merck). Flp-InTM HeLa cells were lentivirally infected with NTC or USP50 shRNA as per the manufacturer‘s protocol and then cells selected using Puromycin. Clones were tested for the ability to knock down expression of FLAG-USP50 after 100 μM IPTG for 72 hours and to phenotypically increase spontaneous 53BP1 foci formation following treatment with USP50 shRNA.

### USP50 gene expression PCR

HeLa cells were plated and transfected with 4 different siRNA sequences for USP50 using Dharmafect. 72 hours later RNA was extracted using the Bioline RNA extraction kit according to the manufacturer’s instructions. RNA was converted into cDNA using the Bioline cDNA synthesis kit. Controls were performed without the reverse transcriptase. USP50 cDNA expression was determined by PCR using the following primers: USP50 Fwd GGAAGTATATCACCGCTCTGC and USP50 Rev TGATCTTCTCCGGGAGTAGTGG. Expression levels of USP50 were normalized to that of GAPDH which was amplified using the following primers: GAPDH_Fwd ATTGTCAGCAATGCATCCTG and GAPDH_Rev ATGGACTGTGGTCATGAGCC

### Plasmid Generation

USP50 was amplified out of the addgene USP50-FLAG plasmid vector (from Wade Harper’s group ^94^) and cloned into pcDNA5/FRT/TO. The pcDNA5/FRT/TO-FLAG-USP50 plasmids were designed and sent to Genscript for synthesis. These plasmids were made siRNA resistant to USP50 siRNA sequences 5 and 7 by introducing a series of silent point mutations as follows: USP50 siRNA sequence 5 - TAT GAT ACC CTT CCA GTT and corresponding siRNA resistant form - TAT GAC ACA CTA CCA GTT A and USP50 siRNA sequence 7 - C TAC CCA GCA TTT ACG and corresponding siRNA resistant form - C TAT CCG GCT TTT ACG.

The pcDNA5/FRT/TO-Myc-FEN1 WT and E359K plasmids were synthesized by Genscript and include siResistance to FEN1 exon 2 siRNA GAUGCCUCUAUGAGCAUUUAU. Likewise, the pcDNA5/FRT/TO-EGFP-WRN WT, E84A and K577M plasmids were synthesized by Genscript and include siResistance to both WRN exon 9 siRNA GAGGGUUUCUAUCUUACUA and WRN exon17 siRNA AUACGUAACUCCAGAAUAC.

The pCW-His-myc-Ubiquitin plasmid was published previously^41^.

### Inducible expression

Flp-InTM HeLa shUSP50 cells were plated in 10 cm2 dishes and transfected with a 4:1 ratio pcDNA5/FRT/TO-FLAG-USP50 variants and the pOG44 Flp Recombinase plasmid using FuGene6. Positive clones were selected with 100 μg/ml hygromycin and tested for expression of FLAG-USP50, by treatment with 2 μg/ml doxycycline for 72 hours and subsequent western blot.

Similarly, Flp-InTM HeLa shUSP50 cells were transfected using FuGene6 with pcDNA5/FRT/TO-Myc-FEN1 variants or pcDNA5/FRT/TO-EGFP-WRN variants and pOG44 in a 4:1 ratio and positive clones selected with hygromycin 100 µg/ml. Expression of inducible genes was confirmed by western blot after incubation with 2 µg/ml doxycycline for 72 hours.

Plasmid and siRNA transfection FuGene6 was used to transfect DNA plasmids into cells at a 3:1 FuGENE (μl) ratio, following the manufacturer’s guidelines. siRNA transfections were carried out using the transfection reagent Dharmafect1 following the manufacturer’s guidelines. For details of siRNA sequences see Supplementary Table 1.

Antibodies Details of antibodies used can be found in **Supplementary Table 2.**

### Colony survival assays

Colony survival assays were used to determine cellular sensitivity in response to hydroxyurea or pyridostatin. Cells were plated at 2 x 105 cells/mL in a 24 well plate and treated with IPTG (100 µM) and / or doxycycline (2 µg/mL) to induce shRNA and FLAG-USP50 expression for 48 hours. Cells were treated with HU (16 hours) or pyridostatin (24 hours) before plating out in a 6 well plate at low density. Plates were incubated for 14 days at 37°C, 5% CO2 until colonies formed. Colonies were stained using 0.5% Crystal violet in 50% methanol and colonies counted.

### Modelling

Molecular graphics and analyses were performed using UCSF ChimeraX, developed by the Resource for Biocomputing, Visualization, and Informatics at the University of California, San Francisco, with support from National Institutes of Health R01-GM129325 and the Office of Cyber Infrastructure and Computational Biology, National Institute of Allergy and Infectious Diseases and Alphafold2 CoLab ^95^. Sequences used were the first Ubiquitin from P0CG47 and USP50 sequence Q70EL3. Note this is not the reference sequence, NP_987090.2, which lacks the sequence “KFLLPS” found in isoform 2.

### Immunofluorescence and microscopy

Cells were plated in a 24 well plate on 13 mm circular glass coverslips at a density of 5 x 104 cells/ml. Cells were treated as required and then fixed in 4% paraformaldehyde (PFA) (unless otherwise stated). Once fixed, cells were permeabilized with 0.2% TritonX100 in PBS, for 5 min, blocked using 10% FBS in PBS for 5 min and incubated with primary antibody for 1 hour at room temperature in 10% FBS/PBS (see Supplementary Table 2 for details). Cells were then washed in 10% FBS/PBS before being incubated for 1 hour with AlexaFluor antibodies at a concentration of 1:2000. Cells were washed in PBS and then fixed for 10 min in 4% PFA before being washed again in PBS. DNA was stained using Hoechst at 1:20,000 for 5 min and then washed with PBS before mounting onto Snowcoat slides using Immunomount mounting media. Cells were imaged on a Leica DM6000B microscope with an HBO lamp with a 100-W mercury short arc UV-bulb light source and four filter cubes, A4, L5, N3 and Y5, to produce excitations at wavelengths of 360, 488, 555 and 647 nm, respectively. Images were captured at each wavelength sequentially with a Plan apochromat HCX 100×/1.4 oil objective at a resolution of 1,392 × 1,040 pixels.

For EdU labelling of S-phase cells, cells were incubated with 10 µM EdU for 10 min prior to fixation. EdU was then labelled with Alexa-fluor-647-azide using Click-IT technology. Briefly, permeablised cells were incubates with the Click-IT reaction cocktail (PBS 1X, 10 uM Biotin Azide, 10 mM Sodium Ascorbate, 1 mM CuSO4) for 30 min at room temperature in the dark. Cells were washed with PBST and labelled with primary and secondary antibodies as above.

### Immunofluorescence and microscopy of ADP-ribosylation

Adherent cells grown on 13 mm circular glass coverslips (Thermo Fisher Scientific) were pre-extracted in 0.2% Triton x-100 in PBS for 2 min on ice, to remove soluble nuclear content, and subsequently fixed with 4% formaldehyde in PBS for 10 min at RT. When cells were stained for PCNA, additional step was added after fixation: coverslips were treated with ice-cold methanol/acetone solution (1:1) for 5 min at room temperature (RT) and washed 3 times for 5 min in PBS. Thereafter, coverslips were blocked with 10% FCS in PBS for 1 h at RT, followed by incubation with appropriate primary antibodies (1 h at RT) and then incubation with fluorochrome-conjugated secondary antibodies (1 h at RT). Coverslips were washed 3 times for 5 min in PBST (PBS with 0.1% Tween20) after both primary and secondary antibody incubations. Next, DNA was stained with DAPI (1 mg/ml in water) for 5 min at RT and coverslips were mounted in fluoroshield (Sigma-Aldrich).

Automated multichannel wide-field microscopy was performed using an Olympus ScanR Screening System equipped with an inverted motorized Olympus IX81 microscope and a motorized stage. Images were acquired using 40x objective at a single autofocus-directed z-position under non-saturating settings. The inbuilt Olympus ScanR Image Analysis Software was used to analyze acquired images. Nuclei were identified by DAPI signal using an integrated intensity-based object detection module. The G1, S and G2 phase cells were gated based on PCNA and DAPI intensity, and fluorescence intensities of interest were quantified.

### Proximity Ligation Assay (PLA)

Flp-InTM HeLa cells were seeded onto poly-L-lysine coated coverslips. For EdU treatment, cells were pulsed with EdU for 10 min (short label) or 24 hours (long label) at 37°C. For analysis of stalled forks, 5 mM HU was added into media following a short EdU pulse for 3 hours at 37°C. Cells were pre-extracted for 5 min on ice with Pre-extraction buffer (20 mM NaCl, 3 mM MgCl2, 300 mM Sucrose, 10 mM PIPES, 0.5% Triton X-100) and fixed in 4% PFA before blocking in 10% BSA overnight. The Click-IT reaction cocktail (PBS 1X, 10 uM Biotin Azide, 10 mM Sodium Ascorbate, 1 mM CuSO4) was added for 1 hour at room temperature in the dark. The Click-IT reaction cocktail was then removed, and cells were incubated in blocking solution for a further 30 min before incubation with primary antibodies (details in Supplementary Table 2) in 10% FBS in PBS for 1 hour at room temperature in the dark. After incubation with primary antibodies, cells were incubated with the corresponding MINUS/PLUS PLA probes (Sigma DUOlink PLA kit) for 1 hour at 37°C in a warm foil-covered box. This was after 2 washes in Buffer A (Sigma DUOlink PLA kit) and then incubated with the PLA kit Ligation solution (1X ligation buffer, ligase enzyme) for 30 min at 37°C. Cells were again washed again in wash buffer A before incubation for 100 min at 37°C with the PLA kit amplification solution (1X amplification buffer, polymerase enzyme). Following amplification cells were washed for 15 min with wash buffer B (Sigma Duolink PLA kit) and incubated with Hoechst for 5 min before another 15 min wash with buffer B. A final 1 min wash in 0.01% wash buffer B was performed. Coverslips were mounted onto glass slides and imaged and quantified the following day using a Leica DM6000 fluorescent microscope with a 100x objective lens.

### FLAG immunoprecipitation

Flp-InTM HeLa FLAG-USP50 cells were plated in a 10 cm plate and treated with doxycycline for 72 hours to express inducible FLAG-USP50. Cells were washed with 10 ml ice-cold 1x PBS before being scraped in ice-cold Nuclear Lysis Buffer (10 mM HEPES pH7.6, 200 mM NaCl, 1.5 mM MgCl2, 10% Glycerol, 0.2 mM EDTA, 1% Triton) for every 10 ml, 1 protease inhibitor tablet (COmplete – SIGMA), 1 phosphatase tablet (PhosSTOP – Roche), 20 μM MG132, 1 μl DNase1 and 200 μl iodoacetamide was added. The lysed cells were then transferred into 1.5 ml Eppendorf tubes and incubated with the nuclear lysis buffer on ice for 1 hour with rotation before centrifugation at 13000 rpm, 4°C for 10 min and the supernatant kept and the pellet discarded. 50 μl of the supernatant was mixed with 20 μl 4x SDS Loading buffer and boiled at 95°C for 5 min. For every IP, 10 μl FLAG-agarose beads were firstly washed out of storage buffer by doing 3x 1ml PBS washes and centrifuging at 3000 rpm between each wash. 60 μl of binding buffer (PBS and nuclear lysis buffer at a ratio of 2:1.5) was added for every 10 μl of agarose beads. The resuspended beads were then added to 450 μl of supernatant for each sample and rotated at 4°C overnight. The following day, the samples were centrifuged at 3000 rpm for 1 min and the beads left to settle. The supernatant was removed before 3x 1 ml PBS-0.02% tween washes. The wash buffer was completely removed before adding 60 μl 2x SDS loading buffer. This was boiled at 95°C for 5 min and 10 μl loaded onto an SDS PAGE gel and analyzed by Western blotting.

### Fibre labelling and spreading

Cells were seeded in 6 cm plates and treated for 72 hours to knock down or overexpress proteins of interest and then treated with thymidine analogues. To monitor on-going replication dynamics, cells were incubated at 37°C with CldU for 20 min at a final concentration of 25 μM and then with CO2-equilibrated IdU at 37°C for 20 min at a final concentration of 250 μM. After incubation with the thymidine analogues, cells were washed once with ice-cold PBS, trypsinized and resuspended in 200 µl of PBS and counted. The optimal final cell density is 50 x 104 cells/ ml and thus cells were adjusted to reach such a concentration. For each sample, three Snowcoat slides were labelled. Near the label of each slide 2 μl of the cell, sample was placed and allowed to slightly dry for 3-4 min. Then 7 μl of spreading buffer (200 mM Tris pH7.4, 50 mM EDTA, 0.5% SDS) was added, mixed with the sample, and incubated for 2 min. To spread the sample down the slide, slides were gradually tilted and once the sample had reached the bottom of the slide, they were allowed to dry for 2 min. Finally, slides were fixed in a 3:1 ratio of Methanol: Acetic acid for 10 min before leaving slides to air dry for 5-10 min. Dried slides were stored at 4°C till staining.

### Fibre Immunostaining

After fibre spreading slides were washed 2 x for 5 min with 1 ml H2O and rinsed with 2.5 M HCl before denaturing DNA with 2.5 M HCl for 1 hour 15 min. Slides were then rinsed 2 x with PBS and washed for 5 min in blocking solution (PBS, 1% BSA, 0.1% Tween20). Slides were incubated for 1 hour in blocking solution. After blocking, each slide was incubated with 115 μl of primary antibodies, Rat αBrdU (Abcam) used at a concentration of 1:1000 and Mouse αBrdU (BD Biosciences) used at 1:750. Slides were covered with large coverslips and incubated with the antibodies for 1 hour. After incubation with the primary antibody, slides were rinsed 3 x with PBS and then incubated for 1 min, 5 min and 25 min, with blocking solution. After rinsing and washing, slides were incubated with 115 μl of secondary antibodies (α-Rat AlexaFluor 555 and α-Mouse AlexaFluor 488) in blocking solution, at a concentration of 1:500, covered with a large coverslip for 2 hours. Slides were rinsed 3 x with PBS and incubated with blocking solution for 1 min, 5 min and 25 min. After again rinsing 2 x with PBS mounting media was added to the slide and a large coverslip placed over the slide and it was left to dry. Coverslips were then stored at -20°C for microscopy analysis. It is important to point out that during this process slides were kept protected from light.

### Fibre Scoring and Analysis

Using 10 image fields of fibres per condition, all visible fibres were assigned a structure (ongoing fork, 1^st^ label origin, 2^nd^ label origin, 1^st^ label termination, 2^nd^ label termination). Then the proportion of each structure was determined based on the total amount of all scored structures containing a first label. For fork asymmetry, only first label origin forks were studied. Both IdU-second label lengths were measured from each side of the CldU first label using the ImageJ software, the lengths compared and given as a proportion of the longest label.

### INDUCE-seq double-strand break mapping and sequence analysis

Cells were harvested in Dulbecco’s phosphate-buffered saline (PBS) and counted. Cells (120,000 per well) were adhered onto a poly-L-lysine–coated 96-well plate and cross-linked in methanol-free paraformaldehyde (final concentration 4%) for 10 min. The PFA was removed, and the wells were washed twice in PBS and stored in 200 μl PBS. Plates were sealed and stored at 4°C until downstream library preparation. INDUCE-seq was performed as previously described ^54^ on Illumina NextSeq500 using 1 × 75 bp high-capacity flow cell. INDUCE-seq was performed in duplicate. After assessing reproducibility by comparing the genome-wide densities of DSBs in 10-kb windows, technical replicates were combined. INDUCE-seq reads were processed as previously described and aligned to the human genome with bowtie2 (GRCh38/hg38)^96^. Using Bedtools ^97^, alignments were converted to Bed files and intersects between biological repeats generated. These were used to generate fasta sequences using Getfasta. Duplicate sequences were removed by Filter Fasta. Nucleotide % were displayed using Fasta Statistics. Oligo-diff was then run comparing the USP50 and NTC data sets to return oligos significantly enriched in one file relative to the other ^98^.

### CO-FISH

HeLa cells were grown in the presence or absence of IPTG (1 mM) to induce shUSP50 expression and in the presence of the telomerase inhibitor BIBR1532 (20 µM) (Cambridge Biosciences CAY16608) for 7 days and COFISH was performed as described previously ^99^. Briefly, cells were cultured in the presence of 2.5 µM BrdU for one population doubling before 0.2 µg/ml colcemid (Gibco 15212012) treatment for the final 4 hours before trypsinization. The cell pellet was resuspended in 75 mM KCl at 37°C for 15 min and fixed in 3:1 methanol:acetic acid and stored at -20°C. Metaphases were dropped onto ice-cold slides and allowed to air dry for 24 hours at room temperature. Slides containing metaphase spreads were treated with RNaseA (100 µg/ml) for 10 min at 37°C, fixed in 4% PFA at room temperature for 10 min and dehydrated in an ethanol series (15%, 85%, 100% ethanol 2 min each) before air-drying. Slides were then incubated with 0.5 µg/ml Hoechst 33258 in 2x saline sodium citrate (SSC) (Sigma) for 15 min and exposed to UV for 30 min in 2x SSC before being treated with exonuclease III (3 U/µl) for a further 10 min. Metaphases were denatured at 72°C in 70% formamide/30% 2xSSC solution for 2 min before hybridization to Alexa 488–OO-(CCCTAA)n probe (Eurogentec) at 37°C for 16 hours. Metaphases were washed 5 times in 2x SSC at 42°C and the DNA counterstained with DAPI (0.1 mg/ml), the slides were then washed in PBS and mounted with immunomount before imaging on a Leica DM6000. 10 metaphases per condition were scored for lost telomeres.

### Chromatin Fractionation

To separate the chromatin-enriched fraction, cells were harvested and washed in PBS, before being re-suspended in sucrose buffer (10 mM Tris-Cl pH 7.5, 20 mM KCl, 250 mM sucrose, 2.5 mM MgCl2, protease inhibitor). Triton X-100 was added to a final concentration of 0.3%, and cells were vortexed 3 x 5 s, followed by centrifugation (500 g, 4°C, 5 min). The supernatant was discarded, and the pellet re-suspended in NETN150 buffer (50 mM Tris-Cl pH 8.0, 150 m NaCl, 2 mM EDTA, 0.5% NP-40, protease inhibitor) and incubated on ice for 30 min, followed by centrifugation (1700 g, 4 °C, 5 min). The supernatant was discarded, and the pellet re-suspended in NETN150 buffer and sonicated 2 x 10 s. Subsequently, 2 x SDS loading buffer was added and the samples were boiled before analysis by Western blotting.

### Statistics

All statistical tests, unless indicated otherwise, used a two-sided unpaired *t*-test: ns = not significant, * = p≤0.05, ** = p≤0.01, *** = p ≤0.001, **** = p≤0.0001. All centre values are given as the mean and all error bars are standard error of the mean (S.E.M.). To aid readability, statistics have only been shown between pertinent groups in figures.

---

**Supplementary Table 1.**
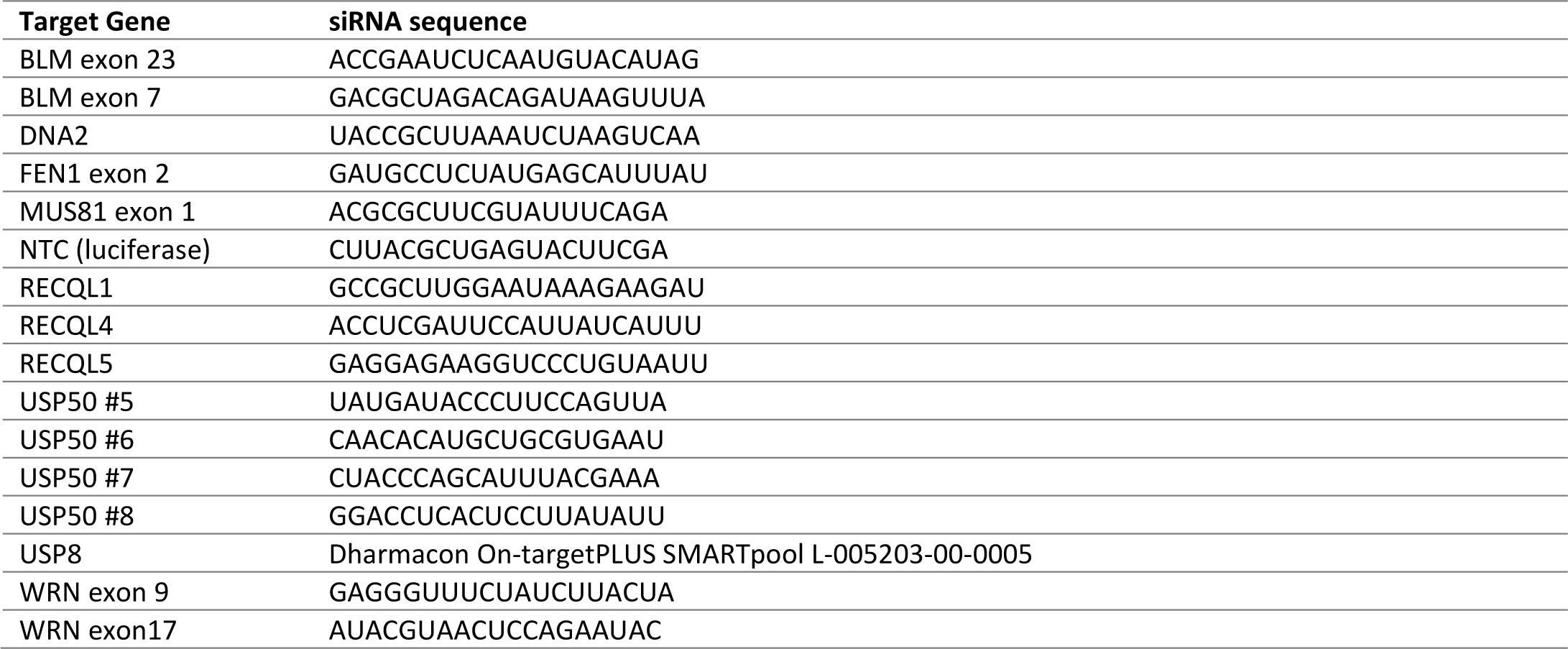
siRNA sequences.

**Supplementary Table 2.** Antibody details.

| Antibody | Animal | Assay | Dilution | Supplier | Cat number | RRID |
| --- | --- | --- | --- | --- | --- | --- |
| 53BP1 | Rabbit | IF | 1:2000 | Abcam | ab36823 | AB_722497 |
| AlexaFluor 647 azide | - | ClickIT | 1:2000 | ThermoFisher | A10277 |  |
| Biotin | Rabbit | PLA | 1:50 | Bethyl | A150-109A | AB_67327 |
| BLM | Goat | WB | 1:1000 | Abcam | ab5446 | AB_304894 |
| BrdU (CldU) | Rat | Fibres | 1:1000 | Abcam | ab6326 | AB_305426 |
| BrdU (IdU) | Mouse | Fibres | 1:750 | BD Biosciences | 347580 | AB_400326 |
| DNA2 | Rabbit | WB/IF | 1:1000/<br>1:200 | Abcam | ab96488 | AB_10677769 |
| Donkey $\alpha$ Mouse AlexaFluor 488 | Donkey | IF | 1:2000 | ThermoFisher | A21202 | AB_141607 |
| Donkey $\alpha$ Mouse AlexaFluor 555 | Goat | IF | 1:2000 | ThermoFisher | A31570 | AB_2536180 |
| Donkey $\alpha$ Rabbit AlexaFluor 488 | Goat | IF | 1:2000 | ThermoFisher | A21206 | AB_2535792 |
| Donkey $\alpha$ Rabbit AlexaFluor555 | Donkey | IF | 1:2000 | ThermoFisher | A31572 | AB_162543 |
| Donkey $\alpha$ Sheep HRP | Donkey | WB | 1:10000 | Sigma | A3415.5ML | AB_258076 |
| Fc-fused anti-poly-ADP-ribose binding reagent | Rabbit | IF |  | Millipore | MABE1031 | AB_2665467 |
| FEN1 | Rabbit | WB/PLA | 1:1000<br>/1:200 | Abcam | ab17994 | AB_444168 |
| FLAG (M2) | Mouse | WB/PLA | 1:2000 /1:300 | Sigma | F1804 | AB_262044 |
| GFP | Mouse | WB | 1:5000 | Roche | 11814460001 | AB_390913 |
| Goat $\alpha$ Rat AlexaFluor 555 | Goat | Fibres | 1:2000 | ThermoFisher | A21434 | AB_141733 |
| Hexa-Histidine | Mouse | WB | 1:1000 | Sigma | H1029 | AB_260015 |
| Histone H2B | Rabbit | WB | 1:1000 | Abcam | ab1790 | AB_302612 |
| Histone H3 | Rabbit | WB | 1:1000 | Abcam | ab1791 | AB_302613 |
| HUS1 | Rabbit | PLA | 1:500 | Proteintech | 11223-1-AP | AB_2248839 |
| Mus81 | Mouse | WB | 1:250 | Novus | NB100-2064 | AB_1109412 |
| Myc | Mouse | WB | 1:2000 | Sigma | M5546 | AB_260581 |
| PCNA | Mouse | PLA | 1:200 | Cell Signaling | 2586 | AB_2160343 |
| Poly/mono-ADP Ribose | Rabbit | IF | 1:1000 | CST | #83732 | AB_2749858 |
| pRPA | Rabbit | WB | 1:1000 | Abcam | ab87277 | AB_1952482 |
| Rabbit $\alpha$ Mouse HRP | Rabbit | WB | 1:5000 | Dako | p0161 | AB_2687969 |
| RECQL1 | Rabbit | WB | 1:3000 | Fisher | PA5-27099 | AB_2544575 |
| RECQL4 | Rabbit | WB | 1:3000 | Proteintech | 17008-1-AP | AB_2238324 |
| RECQL5 | Mouse | WB | 1:1000 | Santa Cruz | 5847 | AB_10834807 |
| RPA | Mouse | WB | 1:3000 | Millipore | NA13 | AB_565121 |
| Swine $\alpha$ Rabbit HRP | Swine | WB | 1:5000 | Dako | P0217 | AB_2728719 |
| Tubulin | Mouse | WB | 1:1000 | Santa Cruz | sc-5286 | AB_628411 |
| USP8 | Sheep | WB | 1:1000 | R&Dsystems | AF7735 | AB_2844099 |
| Vinculin | Rabbit | WB | 1:1000 | Abcam | ab129002 | AB_11144129 |
| WRN | Rabbit | WB/PLA | 1:1000/<br>1:200 | Abcam | ab124673 | AB_10972871 |
| β-actin | Rabbit | WB | 1:2000 | Abcam | ab115777 | AB_10899528 |

**Supplementary Table 3.** – Key Chemicals.

| Reagent | Source | Identifier |
| --- | --- | --- |
| C5 DNA2i | Cambridge Biosciences | HY-128729-5mg |
| FEN1 inhibitor | Synthesized as described in <sup>100</sup>. | Compound 24 in <sup>100</sup>, Compound 1 in <sup>101</sup>. |
| Hydroxyurea | Sigma Aldrich | H8627 |
| Mirin | Alfa Aesar | J67462 |
| PARG inhibitor (PDD 0017273) | Tocris/Sigma Aldrich | 5952/SML1781 |
| PARP inhibitor (Olaparib, AZD2281, Ku-0059436) | ApexBio | A4154 |
| Pyridostatin hydrochloride | Sigma Aldrich | SML2690 |
| Telomerase inhibitor BIBR1532 | Cambridge Biosciences | CAY16608 |

## REFERENCES

1. Zeman MK, Cimprich KA. Causes and consequences of replication stress. Nat Cell Biol 16, 2–9 (2014).

2. Gaillard H, Garcia-Muse T, Aguilera A. Replication stress and cancer. Nat Rev Cancer 15, 276–289 (2015).

3. Lu H, Davis AJ. Human RecQ Helicases in DNA Double-Strand Break Repair. Front Cell Dev Biol 9, 640755 (2021).

4. Larsen NB, Hickson ID. RecQ Helicases: Conserved Guardians of Genomic Integrity. Adv Exp Med Biol 767, 161–184 (2013).

5. Vindigni A, Hickson ID. RecQ helicases: multiple structures for multiple functions? HFSP J 3, 153–164 (2009).

6. Capp C, Wu J, Hsieh TS. RecQ4: the second replicative helicase? Crit Rev Biochem Mol Biol 45, 233–242 (2010).

7. Abu-Libdeh B, et al. RECON syndrome is a genome instability disorder caused by mutations in the DNA helicase RECQL1. J Clin Invest 132, (2022).

8. Mojumdar A. Mutations in conserved functional domains of human RecQ helicases are associated with diseases and cancer: A review. Biophys Chem 265, 106433 (2020).

9. Chan EM, et al. WRN helicase is a synthetic lethal target in microsatellite unstable cancers. Nature 568, 551–556 (2019).

10. Lieb S, et al. Werner syndrome helicase is a selective vulnerability of microsatellite instability-high tumor cells. Elife 8, (2019).

11. Kategaya L, Perumal SK, Hager JH, Belmont LD. Werner Syndrome Helicase Is Required for the Survival of Cancer Cells with Microsatellite Instability. iScience 13, 488–497 (2019).

12. Behan FM, et al. Prioritization of cancer therapeutic targets using CRISPR-Cas9 screens. Nature 568, 511–516 (2019).

13. van Wietmarschen N, et al. Repeat expansions confer WRN dependence in microsatellite-unstable cancers. Nature 586, 292–298 (2020).

14. Davies SL, North PS, Hickson ID. Role for BLM in replication-fork restart and suppression of origin firing after replicative stress. Nat Struct Mol Biol 14, 677–679 (2007).

15. Berti M, et al. Human RECQ1 promotes restart of replication forks reversed by DNA topoisomerase I inhibition. Nat Struct Mol Biol 20, 347–354 (2013).

16. Zellweger R, et al. Rad51-mediated replication fork reversal is a global response to genotoxic treatments in human cells. J Cell Biol 208, 563–579 (2015).

17. Rodriguez-Lopez AM, Jackson DA, Iborra F, Cox LS. Asymmetry of DNA replication fork progression in Werner’s syndrome. Aging Cell 1, 30–39 (2002).

18. Sidorova JM, Li N, Folch A, Monnat RJ, Jr. The RecQ helicase WRN is required for normal replication fork progression after DNA damage or replication fork arrest. Cell Cycle 7, 796–807 (2008).

19. Su F, et al. Nonenzymatic Role for WRN in Preserving Nascent DNA Strands after Replication Stress. Cell Reports 9, 1387–1401 (2014).

20. Baynton K, Otterlei M, Bjoras M, von Kobbe C, Bohr VA, Seeberg E. WRN interacts physically and functionally with the recombination mediator protein RAD52. J Biol Chem 278, 36476–36486 (2003).

21. Pichierri P, Nicolai S, Cignolo L, Bignami M, Franchitto A. The RAD9-RAD1-HUS1 (9.1.1) complex interacts with WRN and is crucial to regulate its response to replication fork stalling. Oncogene 31, 2809–2823 (2012).

22. Thangavel S, et al. DNA2 drives processing and restart of reversed replication forks in human cells. J Cell Biol 208, 545–562 (2015).

23. Blundred R, Myers K, Helleday T, Goldman AS, Bryant HE. Human RECQL5 overcomes thymidine-induced replication stress. DNA Repair (Amst) 9, 964–975 (2010).

24. Popuri V, Huang J, Ramamoorthy M, Tadokoro T, Croteau DL, Bohr VA. RECQL5 plays co-operative and complementary roles with WRN syndrome helicase. Nucleic Acids Res 41, 881–899 (2013).

25. Kanagaraj R, Saydam N, Garcia PL, Zheng L, Janscak P. Human RECQ5beta helicase promotes strand exchange on synthetic DNA structures resembling a stalled replication fork. Nucleic Acids Res 34, 5217–5231 (2006).

26. Ozsoy AZ, Ragonese HM, Matson SW. Analysis of helicase activity and substrate specificity of Drosophila RECQ5. Nucleic Acids Res 31, 1554–1564 (2003).

27. Rossi ML, Ghosh AK, Bohr VA. Roles of Werner syndrome protein in protection of genome integrity. DNA Repair (Amst) 9, 331–344 (2010).

28. Rogers CM, Wang JC, Noguchi H, Imasaki T, Takagi Y, Bochman ML. Yeast Hrq1 shares structural and functional homology with the disease-linked human RecQ4 helicase. Nucleic Acids Res 45, 5217–5230 (2017).

29. Hudson JJR, Rass U. DNA2 in Chromosome Stability and Cell Survival-Is It All about Replication Forks? Int J Mol Sci 22, (2021).

30. Peng G, et al. Human nuclease/helicase DNA2 alleviates replication stress by promoting DNA end resection. Cancer Res 72, 2802–2813 (2012).

31. Pinto C, Kasaciunaite K, Seidel R, Cejka P. Human DNA2 possesses a cryptic DNA unwinding activity that functionally integrates with BLM or WRN helicases. Elife 5, (2016).

32. Sturzenegger A, et al. DNA2 cooperates with the WRN and BLM RecQ helicases to mediate long-range DNA end resection in human cells. J Biol Chem 289, 27314–27326 (2014).

33. Duxin JP, et al. Okazaki fragment processing-independent role for human Dna2 enzyme during DNA replication. J Biol Chem 287, 21980–21991 (2012).

34. Duxin JP, et al. Human Dna2 is a nuclear and mitochondrial DNA maintenance protein. Mol Cell Biol 29, 4274–4282 (2009).

35. Li Z, et al. hDNA2 nuclease/helicase promotes centromeric DNA replication and genome stability. EMBO J 37, (2018).

36. Shaheen R, et al. Genomic analysis of primordial dwarfism reveals novel disease genes. Genome Res 24, 291–299 (2014).

37. Tarnauskaite Z, et al. Biallelic variants in DNA2 cause microcephalic primordial dwarfism. Hum Mutat 40, 1063–1070 (2019).

38. Di Lazzaro Filho R, et al. Biallelic variants in DNA2 cause poikiloderma with congenital cataracts and severe growth failure reminiscent of Rothmund-Thomson syndrome. J Med Genet, (2023).

39. Ulrich HD. Two-way communications between ubiquitin-like modifiers and DNA. Nat Struct Mol Biol 21, 317–324 (2014).

40. Smeenk G, Mailand N. Writers, Readers, and Erasers of Histone Ubiquitylation in DNA Double-Strand Break Repair. Front Genet 7, 122 (2016).

41. Butler LR, et al. The proteasomal de-ubiquitinating enzyme POH1 promotes the double-strand DNA break response. The EMBO journal 31, 3918–3934 (2012).

42. Yuan J, et al. HERC2-USP20 axis regulates DNA damage checkpoint through Claspin. Nucleic Acids Res 42, 13110–13121 (2014).

43. Quesada Vc, Dıá z-Perales A, Gutiérrez-Fernández A, Garabaya C, Cal S, López-Otıń C. Cloning and enzymatic analysis of 22 novel human ubiquitin-specific proteases. Biochemical and Biophysical Research Communications 314, 54–62 (2004).

44. Ernst A, et al. A strategy for modulation of enzymes in the ubiquitin system. Science 339, 590–595 (2013).

45. Komander D, Clague MJ, Urbe S. Breaking the chains: structure and function of the deubiquitinases. Nat Rev Mol Cell Biol 10, 550–563 (2009).

46. Dantuma NP, Acs K, Luijsterburg MS. Should I stay or should I go: VCP/p97-mediated chromatin extraction in the DNA damage response. Exp Cell Res 329, 9–17 (2014).

47. Petermann E, Luis Orta M, Issaeva N, Schultz N, Helleday T. Hydroxyurea-Stalled Replication Forks Become Progressively Inactivated and Require Two Different RAD51-Mediated Pathways for Restart and Repair. Molecular Cell 37, 492–502 (2010).

48. Karlsson M, et al. A single-cell type transcriptomics map of human tissues. Sci Adv 7, (2021).

49. Byun TS, Pacek M, Yee MC, Walter JC, Cimprich KA. Functional uncoupling of MCM helicase and DNA polymerase activities activates the ATR-dependent checkpoint. Genes Dev 19, 1040–1052 (2005).

50. Nickoloff JA, Sharma N, Taylor L, Allen SJ, Hromas R. The Safe Path at the Fork: Ensuring Replication-Associated DNA Double-Strand Breaks are Repaired by Homologous Recombination. Front Genet 12, 748033 (2021).

51. Hanada K, et al. The structure-specific endonuclease Mus81 contributes to replication restart by generating double-strand DNA breaks. Nature Structural & Molecular Biology 14, 1096–1104 (2007).

52. Franchitto A, Pirzio LM, Prosperi E, Sapora O, Bignami M, Pichierri P. Replication fork stalling in WRN-deficient cells is overcome by prompt activation of a MUS81-dependent pathway. J Cell Biol 183, 241–252 (2008).

53. Rass U. Resolving branched DNA intermediates with structure-specific nucleases during replication in eukaryotes. Chromosoma 122, 499–515 (2013).

54. Dobbs FM, van Eijk P, Fellows MD, Loiacono L, Nitsch R, Reed SH. Precision digital mapping of endogenous and induced genomic DNA breaks by INDUCE-seq. Nat Commun 13, 3989 (2022).

55. Piovesan A, Pelleri MC, Antonaros F, Strippoli P, Caracausi M, Vitale L. On the length, weight and GC content of the human genome. BMC Res Notes 12, 106 (2019).

56. Li B, Reddy S, Comai L. The Werner Syndrome Helicase Coordinates Sequential Strand Displacement and FEN1-Mediated Flap Cleavage during Polymerase delta Elongation. Mol Cell Biol 37, (2017).

57. Sarkies P, Murat P, Phillips LG, Patel KJ, Balasubramanian S, Sale JE. FANCJ coordinates two pathways that maintain epigenetic stability at G-quadruplex DNA. Nucleic Acids Res 40, 1485–1498 (2012).

58. Sharma S, et al. WRN helicase and FEN-1 form a complex upon replication arrest and together process branchmigrating DNA structures associated with the replication fork. Mol Biol Cell 15, 734–750 (2004).

59. Brosh RM, Jr., et al. Werner syndrome protein interacts with human flap endonuclease 1 and stimulates its cleavage activity. EMBO J 20, 5791–5801 (2001).

60. Saharia A, Teasley DC, Duxin JP, Dao B, Chiappinelli KB, Stewart SA. FEN1 ensures telomere stability by facilitating replication fork re-initiation. J Biol Chem 285, 27057–27066 (2010).

61. Wang W, et al. The human Rad9-Rad1-Hus1 checkpoint complex stimulates flap endonuclease 1. Proc Natl Acad Sci U S A 101, 16762–16767 (2004).

62. Iannascoli C, Palermo V, Murfuni I, Franchitto A, Pichierri P. The WRN exonuclease domain protects nascent strands from pathological MRE11/EXO1-dependent degradation. Nucleic Acids Res 43, 9788–9803 (2015).

63. Chung L, et al. The FEN1 E359K germline mutation disrupts the FEN1-WRN interaction and FEN1 GEN activity, causing aneuploidy-associated cancers. Oncogene 34, 902–911 (2015).

64. Hanzlikova H, Kalasova I, Demin AA, Pennicott LE, Cihlarova Z, Caldecott KW. The Importance of Poly(ADP-Ribose) Polymerase as a Sensor of Unligated Okazaki Fragments during DNA Replication. Mol Cell 71, 319–331 e313 (2018).

65. Crabbe L, Verdun RE, Haggblom CI, Karlseder J. Defective telomere lagging strand synthesis in cells lacking WRN helicase activity. Science 306, 1951–1953 (2004).

66. Saharia A, et al. Flap endonuclease 1 contributes to telomere stability. Curr Biol 18, 496–500 (2008).

67. Damerla RR, Knickelbein KE, Strutt S, Liu FJ, Wang H, Opresko PL. Werner syndrome protein suppresses the formation of large deletions during the replication of human telomeric sequences. Cell Cycle 11, 3036–3044 (2012).

68. Liu W, et al. A Selective Small Molecule DNA2 Inhibitor for Sensitization of Human Cancer Cells to Chemotherapy. EBioMedicine 6, 73–86 (2016).

69. Bryant HE, et al. PARP is activated at stalled forks to mediate Mre11-dependent replication restart and recombination. Embo J 28, 2601–2615 (2009).

70. Nimonkar AV, et al. BLM-DNA2-RPA-MRN and EXO1-BLM-RPA-MRN constitute two DNA end resection machineries for human DNA break repair. Genes Dev 25, 350–362 (2011).

71. Aressy B, et al. A screen for deubiquitinating enzymes involved in the G(2)/M checkpoint identifies USP50 as a regulator of HSP90-dependent Wee1 stability. Cell Cycle 9, 3815–3822 (2010).

72. Cai J, Wei J, Schrott V, Zhao J, Bullock G, Zhao Y. Induction of deubiquitinating enzyme USP50 during erythropoiesis and its potential role in the regulation of Ku70 stability. J Investig Med 66, 1–6 (2018).

73. Lee JY, et al. The deubiquitinating enzyme, ubiquitin-specific peptidase 50, regulates inflammasome activation by targeting the ASC adaptor protein. FEBS Lett 591, 479–490 (2017).

74. Buus R, Faronato M, Hammond DE, Urbe S, Clague MJ. Deubiquitinase activities required for hepatocyte growth factor-induced scattering of epithelial cells. Curr Biol 19, 1463–1466 (2009).

75. Groza T, et al. The International Mouse Phenotyping Consortium: comprehensive knockout phenotyping underpinning the study of human disease. Nucleic Acids Res 51, D1038–D1045 (2023).

76. Gray MD, et al. The Werner syndrome protein is a DNA helicase. Nature Genetics 17, 100–103 (1997).

77. Brosh RM, Jr., Opresko PL, Bohr VA. Enzymatic mechanism of the WRN helicase/nuclease. Methods Enzymol 409, 52–85 (2006).

78. Hansel-Hertsch R, Spiegel J, Marsico G, Tannahill D, Balasubramanian S. Genome-wide mapping of endogenous G-quadruplex DNA structures by chromatin immunoprecipitation and high-throughput sequencing. Nat Protoc 13, 551–564 (2018).

79. Lyu J, Shao R, Kwong Yung PY, Elsasser SJ. Genome-wide mapping of G-quadruplex structures with CUT&Tag. Nucleic Acids Res 50, e13 (2022).

80. Marsico G, et al. Whole genome experimental maps of DNA G-quadruplexes in multiple species. Nucleic Acids Res 47, 3862–3874 (2019).

81. Speina E, et al. Human RECQL5beta stimulates flap endonuclease 1. Nucleic Acids Res 38, 2904–2916 (2010).

82. Rossi ML, Ghosh AK, Kulikowicz T, Croteau DL, Bohr VA. Conserved helicase domain of human RecQ4 is required for strand annealing-independent DNA unwinding. DNA Repair (Amst) 9, 796–804 (2010).

83. Budhathoki JB, Ray S, Urban V, Janscak P, Yodh JG, Balci H. RecQ-core of BLM unfolds telomeric G-quadruplex in the absence of ATP. Nucleic Acids Res 42, 11528–11545 (2014).

84. Thangavel S, et al. Human RECQ1 and RECQ4 helicases play distinct roles in DNA replication initiation. Mol Cell Biol 30, 1382–1396 (2010).

85. Xu X, Rochette PJ, Feyissa EA, Su TV, Liu Y. MCM10 mediates RECQ4 association with MCM2-7 helicase complex during DNA replication. EMBO J 28, 3005–3014 (2009).

86. Hu Y, et al. RECQL5/Recql5 helicase regulates homologous recombination and suppresses tumor formation via disruption of Rad51 presynaptic filaments. Genes Dev 21, 3073–3084 (2007).

87. Saponaro M, et al. RECQL5 controls transcript elongation and suppresses genome instability associated with transcription stress. Cell 157, 1037–1049 (2014).

88. Walden M, Masandi SK, Pawlowski K, Zeqiraj E. Pseudo-DUBs as allosteric activators and molecular scaffolds of protein complexes. Biochem Soc Trans 46, 453–466 (2018).

89. Morrow ME, et al. Active site alanine mutations convert deubiquitinases into high-affinity ubiquitin-binding proteins. EMBO Rep 19, (2018).

90. Lee JG, Baek K, Soetandyo N, Ye Y. Reversible inactivation of deubiquitinases by reactive oxygen species in vitro and in cells. Nat Commun 4, 1568 (2013).

91. Mirsanaye AS, Typas D, Mailand N. Ubiquitylation at Stressed Replication Forks: Mechanisms and Functions. Trends Cell Biol 31, 584–597 (2021).

92. Dungrawala H, Cortez D. Purification of proteins on newly synthesized DNA using iPOND. Methods Mol Biol 1228, 123–131 (2015).

93. Wessel SR, Mohni KN, Luzwick JW, Dungrawala H, Cortez D. Functional Analysis of the Replication Fork Proteome Identifies BET Proteins as PCNA Regulators. Cell Rep 28, 3497–3509 e3494 (2019).

94. Sowa ME, Bennett EJ, Gygi SP, Harper JW. Defining the human deubiquitinating enzyme interaction landscape. Cell 138, 389–403 (2009).

95. Jumper J, et al. Highly accurate protein structure prediction with AlphaFold. Nature 596, 583–589 (2021).

96. Langmead B, Salzberg SL. Fast gapped-read alignment with Bowtie 2. Nat Methods 9, 357–359 (2012).

97. Quinlan AR, Hall IM. BEDTools: a flexible suite of utilities for comparing genomic features. Bioinformatics 26, 841–842 (2010).

98. Santana-Garcia W, et al. RSAT 2022: regulatory sequence analysis tools. Nucleic Acids Res 50, W670–W676 (2022).

99. Bailey SM, Cornforth MN, Kurimasa A, Chen DJ, Goodwin EH. Strand-specific postreplicative processing of mammalian telomeres. Science 293, 2462–2465 (2001).

100. Tumey LN, et al. The identification and optimization of a N-hydroxy urea series of flap endonuclease 1 inhibitors. Bioorg Med Chem Lett 15, 277–281 (2005).

101. Exell JC, et al. Cellularly active N-hydroxyurea FEN1 inhibitors block substrate entry to the active site. Nat Chem Biol 12, 815–821 (2016).

